# Reversal effects of Isochlorogenic acid A on HBV-induced transcriptional dysregulation and apoptotic signaling

**DOI:** 10.64898/2026.06.23.733975

**Authors:** Giscard Wilfried Koyaweda, Mirco Glitscher, Csaba Miskey, Eberhard Hildt

## Abstract

Chronic hepatitis B virus (HBV) infection contributes to hepatocellular carcinoma by disrupting host transcription, cell-cycle control, and apoptotic signaling. Isochlorogenic acid A (ICAA), a natural compound with antiviral and hepatoprotective properties, was previously shown to inhibit HBV replication by interfering with multiple steps of the viral life cycle. Because chronic HBV often reflects an imbalance between proliferation and cell death, we investigated how ICAA affects gene expression related to these processes in the presence or absence of HBV.

We performed transcriptome analysis using RNA sequencing (RNA-seq) in HepAD38 cells (a HepG2-derived stable HBV-expressing line) and HepG2 control cells (HBV-negative) treated with ICAA or DMSO. HBV caused major differences in gene expression in HepAD38 cells compared with HBV-negative HepG2 cells. Principal component analysis showed that ICAA significantly altered HBV-dependent expression patterns, resulting in 189 differentially expressed genes (DEGs) that were regulated in opposite directions by both HBV and ICAA.

Functional enrichment analysis highlighted pathways in viral carcinogenesis, apoptosis, MAPK signaling, and p53 signaling. Annexin V/propidium iodide assays showed apoptotic cells in both treated and untreated HepAD38 cultures, with only minor pattern changes. Mechanistically, in untreated HBV-positive cells caspase-9 cleavage failed to activate PARP, suggesting that induction of intrinsic apoptosis is followed by blocked execution. In contrast, ICAA inhibits caspase-9 cleavage in a dose-dependent manner, while activating PARP. Consistent with this, ICAA treatment increased apoptotic DNA fragmentation in HepAD38, reflecting the proapoptotic potential of ICAA under these conditions facilitating the elimination of HBV-positive cells by apoptosis. These findings highlight the potential therapeutic relevance of this compound in processes associated with HBV pathogenesis, together with its antiviral effect.

**Graphical abstract:** 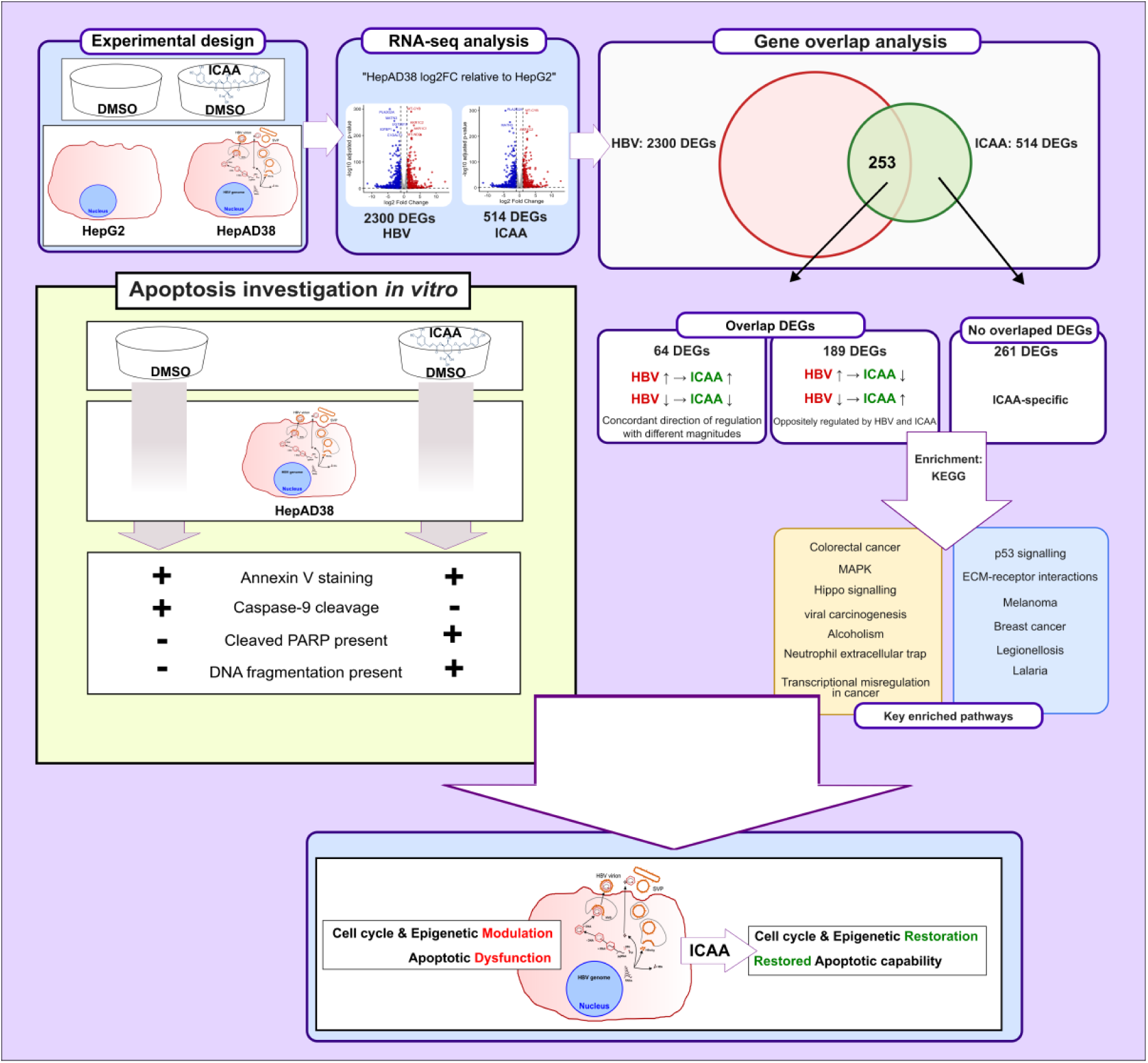

## Introduction

Hepatitis B virus (HBV) is a small DNA virus that almost exclusively infects human hepatocytes. HBV harbors a partially double-stranded DNA genome that encompasses a ∼3.2 kb genome and encodes four overlapping genes: S (surface proteins), C (core protein, precore precursor of HBeAg), P (polymerase), and X (regulatory HBx protein) [1, 2]. Upon entry, the relaxed circular DNA (rcDNA) is converted to nuclear covalently closed circular DNA (cccDNA), the reservoir responsible for persistence and treatment challenge [3–5]. Despite the existence of a highly effective vaccine [6] the global burden of chronic hepatitis B virus (HBV) infection is estimated to be around 254 million, resulting in significant morbidity from cirrhosis and hepatocellular carcinoma. Curing chronic HBV infection remains a major challenge.

Current treatments involving interferons (IFNs) and nucleos(t)ide analogues (NAs) can suppress HBV, but they cannot eliminate the virus because the virus’s cccDNA remains in the nucleus of infected cells. These treatments clearly reduce the risk of cirrhosis, hepatic failure and hepatocellular carcinoma (HCC), but require long-term administration. [7]. In addition to the high stability of the cccDNA, integration of HBV DNA and the lack of an immune response are other obstacles to the treatment [8]. HBV proteins, HBx in particular, interfere with the regulation of the host cell cycle, induce genomic instability and modulate apoptosis. These are key events that facilitate virus persistence and allow hepatocarcinogenesis. While the effect of HBV on apoptosis is a subject of conflicting findings in the literature, several studies have demonstrated that HBV alters apoptotic signaling, thereby favoring viral persistence and contributing to liver injury and hepatocellular carcinoma development [9, 10]. The interplay between viral replication, viral proteins interacting with host cells, and the resulting progression of hepatic disease underscores the necessity of novel therapeutic strategies that target both HBV and host mechanisms directly. [11].

Ischlorogenic acid A (ICAA), also known as 3,5-dicaffeoylquinic acid, is a natural polyphenol derived from caffeic and quinic acids, which has antiviral activity against a variety of viruses [12–14]. We have already described its antiviral activity against HBV, through interference with several stages of the viral replication cycle [14]. In addition to its antiviral effects, ICAA has also been shown to exert hepatoprotective and neuroprotective effects as well as antioxidant properties [15–17]. Given the association between chronic HBV infection with cirrhosis, hepatic failure, and hepatocellular carcinoma (HCC), we performed a transcriptomic analysis to elucidate the effect of ICAA on HBV-associated pathogenesis. Our data show that several transcripts, including key genes involved in HBV-related disease progression, are regulated in opposite directions by both ICAA and HBV. Further in vitro analyses suggest that ICAA selectively reverses HBV-dependent metabolic and survival adaptations. Moreover, HBV-dependent impaired apoptosis is rescued, leading to execution of cell death in HBV-expressing HepAD38. These data provide new molecular insight into how ICAA can contribute to suppressing the development of HBV-induced hepatocellular carcinoma.

## 1. Methodology

### 1.1. Cell culture and treatments

HepG2 cells were maintained in Roswell Park Memorial Institute 1640 (RPMI 1640) medium supplemented with 10% (v/v) fetal bovine serum (FBS) (FBS.S 0615, Bio & Sell GmbH), 2 mM L-glutamine (Q), 100 μg/mL streptomycin (S), and 100 U/mL penicillin (P). HepAD38 cells [18] were grown in Dulbecco’s Modified Eagle Medium (DMEM) supplemented with 10% (v/v) FBS, 2 mM Q, 100 μg/mL S, 100 U/mL P, 50 mM hydrocortisone 21-hemisuccinate (sc-250130, Santa Cruz), and 5 μg/mL insulin (I6634-100MG, Sigma-Aldrich). In the HepAD38 culture medium, tetracycline was consistently absent.

For RNA analysis, cells were seeded in 6-well plates at a density of 5 x 10^5^ and treated with 200 μM ICAA for 1 or 3 days, as previously demonstrated to be effective against HBV without cytotoxicity [14]. For downstream analysis, concentrations of 0-500 μM ICAA were used to determine the dose-dependent effects.

### 1.2. High-throughput RNA sequencing (RNA-Seq)

Total RNA was isolated from samples collected at the indicated post-treatment time points using the Direct-zol RNA Miniprep Kit (Zymo Research). RNA integrity was assessed on a Fragment Analyzer (Agilent), and only preparations with an RNA Quality Number (RQN) greater than 9 were selected for downstream analyses. Poly(A)+ RNA was enriched from 1 μg of total RNA using the NEBNext Poly(A) mRNA Magnetic Isolation Module (New England Biolabs).

Sequencing libraries were generated using a modified non-self-complementary random (NNSR) priming strategy as previously described [19]. For first-strand cDNA synthesis, 5 μL of purified mRNA was combined with 100 pmol *NNSR_RT* primer (**Table 1**), denatured at 65°C for 5 min, and immediately cooled on ice. Reverse transcription was carried out in the presence of first-strand synthesis buffer, dithiothreitol, dNTPs, RiboLock RNase inhibitor, and Maxima Reverse Transcriptase (Thermo Fisher Scientific) in a final volume of 20 μL. Reactions were incubated at 45°C for 30 min, followed by enzyme inactivation at 70°C for 15 min.

**Table 1.**
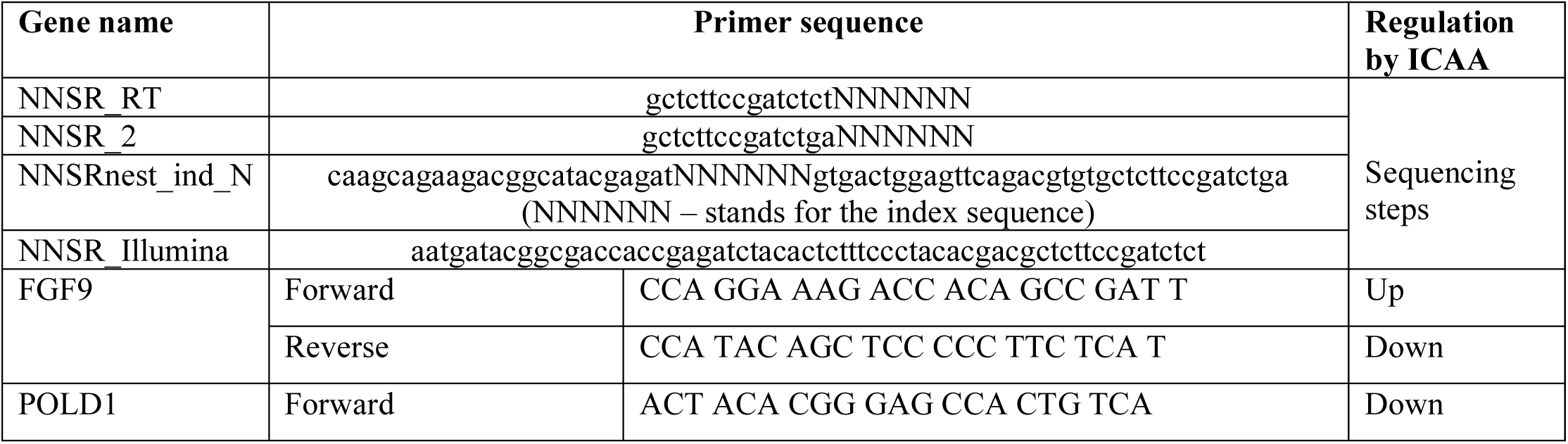
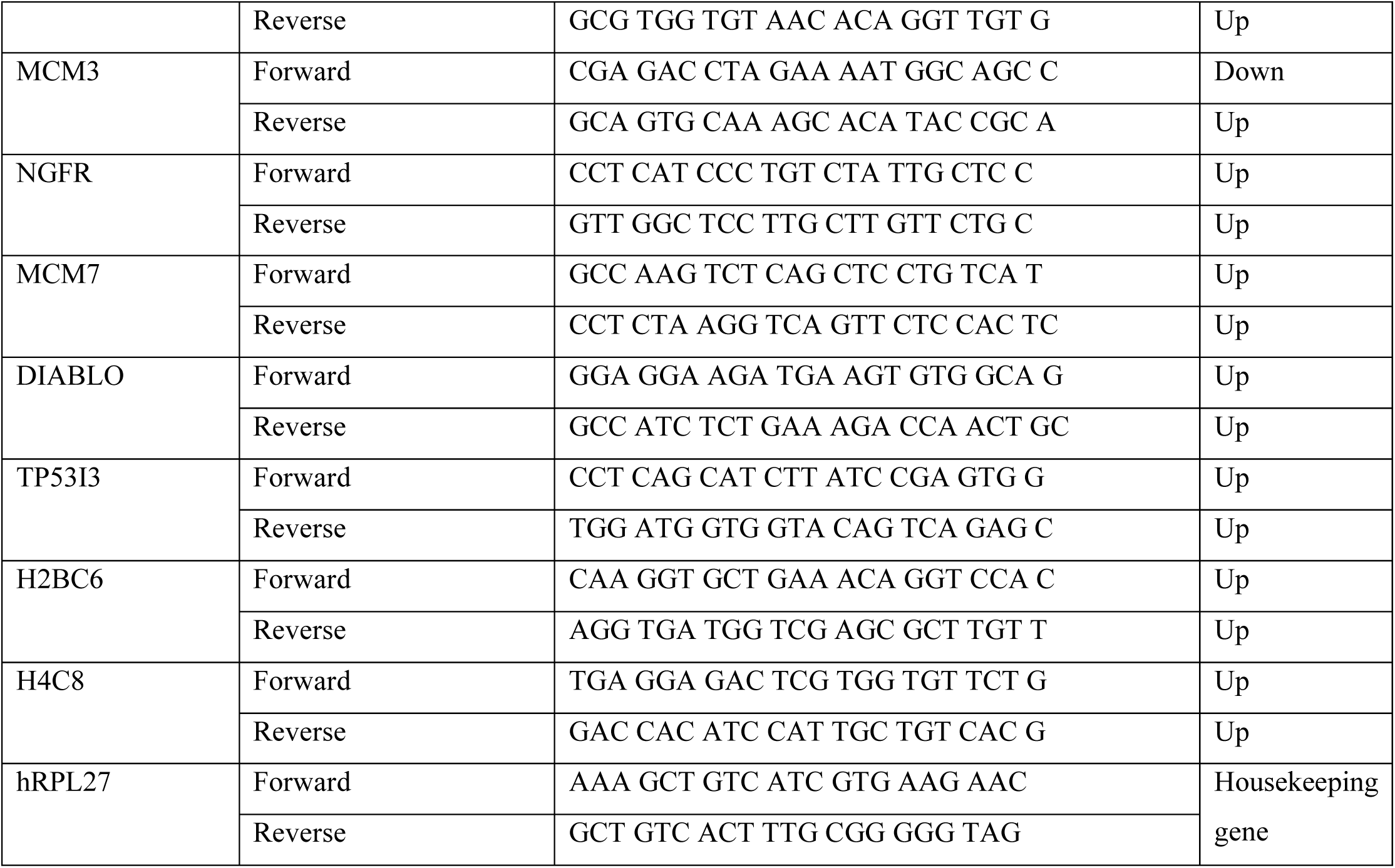
List of primers used in the present study.

Following magnetic bead purification, residual RNA was removed by RNase H digestion at 37°C for 20 min. The resulting first-strand cDNA was purified and converted into double-stranded cDNA using 3′–5′ exonuclease-deficient Klenow Fragment (NEB) and the *NNSR-2* primer (**Table 1**). Second-strand synthesis reactions were performed at 37°C for 30 min in the presence of NEB buffer 2 and dNTPs.

After purification, double-stranded cDNA was amplified and indexed by PCR using NEBNext Ultra II Master Mix (NEB) together with the NNSRnest_ind_N and NNSR_Illumina primers (**Table 1**). Amplification consisted of an initial denaturation step, followed by five cycles at a lower annealing temperature and fifteen cycles at a higher annealing temperature to generate barcoded Illumina-compatible libraries.

PCR products corresponding to approximately 300–500 bp were size-selected from 1.5% agarose gels and recovered using the Zymoclean Gel DNA Recovery Kit (Zymo Research). Libraries were quantified by qPCR with the NEBNext Library Quantification Kit for Illumina and sequenced on an Illumina NextSeq 2000 platform using a single-end 130-cycle configuration.

### 1.3. Reads Processing and differential gene expression analysis

Raw sequencing reads were subjected to adapter removal and quality filtering using *fastp* [20]. Filtered reads were subsequently aligned to the human reference genome assembly hg38 using *STAR [21]* with default alignment parameters. Differential gene expression analysis was performed in R using the *DESeq2* package [22], following the Bioconductor RNA-seq workflow recommendations (https://master.bioconductor.org/packages/release/workflows/vignettes/rnaseqGene/inst/doc/rnaseqGene.html). Functional enrichment analyses were conducted using Kyoto Encyclopedia of Genes and Genomes (KEGG) pathway annotations [23]. Pathway enrichment and visualization were carried out in *R* version 4.4 (https://www.R-project.org) with the *clusterProfiler* [24] and *pathview* packages.

### 1.4. RNA-seq results validation

For *quantitative* Polymerase Chain Reaction (qPCR) analysis, RNA was reverse transcribed to complementary DNA (cDNA) by using the RevertAid H minus RT Kit (Thermo Scientific, EP0452). Randomly selected genes with their corresponding primers as shown in **Table 1** were used for validation of RNA-seq data by qPCR using the GreenMasterMix (Genaxxon; Cat# M3023).

### 1.5. Apoptosis detection

Annexin V staining was performed using the Annexin V-FITC Kit (Militenyi Biotech, Order no. 130-092-052) and analyzed with the MACSQuant Analyzer 10. Apoptotic populations were quantified using FlowJo software (Version 10).

For Western blot experiments, samples were denatured in sample loading buffer at 95 °C for 10 min and then separated by sodium dodecyl sulfate-polyacrylamide gel electrophoresis (SDS-PAGE). The proteins were transferred onto a polyvinylidene difluoride membrane (PVDF) (P667.1, Carl Roth). For detection of the blotted proteins, the following primary antibodies were used: cleaved PARP (Asp214) and caspase-9 (9546 and 9502, respectively; Cell Signaling), and β-Actin (A5316; Sigma).

TUNEL assay was performed using DeadEnd™ Fluorometric TUNEL System (Promega) according to the manufacturer’s procedure.

### 1.6. Statistical Analysis

Data analysis was performed using GraphPad Prism software (v.10.6.1, GraphPad). Data sets presented as fold-change values were calculated relative to the respective control sample in each independent experiment, which was arbitrarily set to 1. For statistical analysis, the data sets were analyzed for significance using an unpaired, two-tailed Student’s t-test. A *p*-value less than 0.05 was considered significant, with significance indicated as follows: \**p* < 0.05, \*\**p* < 0.01, \*\*\**p* < 0.001, \*\*\*\**p* < 0.0001.

## 2. Results

### 2.1. Identification and analysis of differentially expressed genes (DEGs)

To investigate the DEGs induced by HBV and ICAA, two cell lines were used: HepG2 (a HBV-negative) and HepAD38 (a HBV-expressing cell line derived from HepG2 cells). Four biological replicates were performed for each condition, with the cells cultivated under the same passages. Cultivation was performed for 1 or 3 days. RNA-seq produced raw read counts at the gene level reflecting the expression level of the respective genes in the samples. To identify the experimental conditions showing the strongest transcriptional separation for subsequent downstream analyses, a principal component analysis (PCA) on the normalized expression profiles was performed to assess the global structure of the dataset across the different cell lines, treatments, and time points. PCA revealed time-dependent transcriptional changes upon ICAA treatment in HBV-expressing cells, with robust change occurring 3 days post-treatment (**Fig. 1**). However, only moderate changes could be observed in HepG2 cells conditions, prompting our main analysis on HepAD38 cells 3 days post-treatment.

**Fig. 1.**
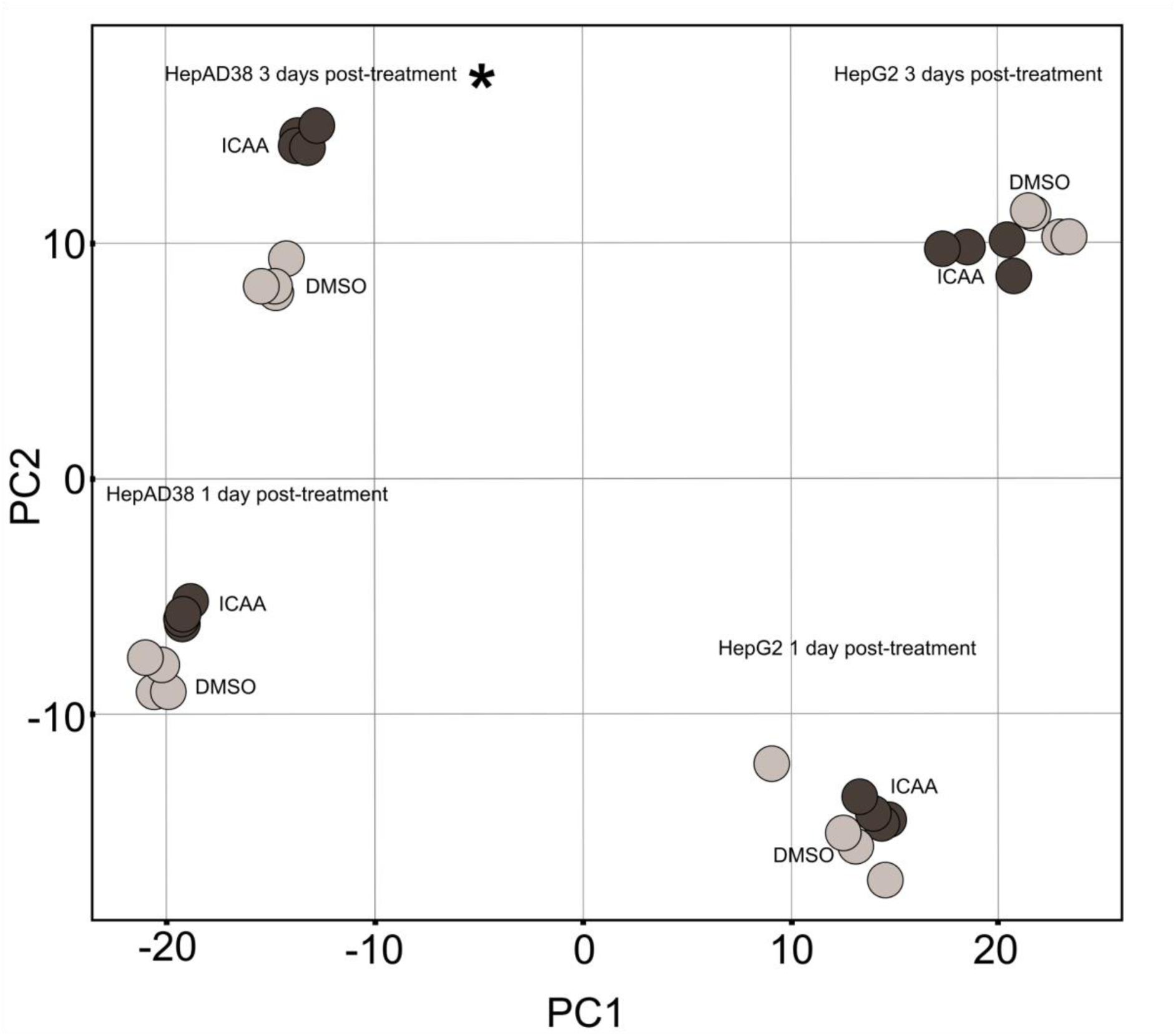
Principal component analysis (PCA) of transcriptomic profiles in HepG2 and HepAD38 cells treated with ICAA or DMSO. PCA was performed using transcriptomic datasets obtained from HepG2 and HepAD38 cells treated with either DMSO or ICAA for 1 or 3 days. Four biological replicates were included for each experimental condition. ICAA-treated samples are represented in dark grey, while DMSO-treated controls are shown in light grey. The cell lines and treatment time points are indicated in the figure.

To determine transcriptomic changes between the different conditions, the log2 fold-change (log2FC) was used as a measure of contrast. DMSO-treated HepG2 cells were used as the baseline control required for these calculations. DEGs analysis of DMSO-treated HepAD38 cells revealed 2,300 HBV-induced DEGs when compared to DMSO-treated HepG2 cells for the three days post-treatment condition, while ICAA treatment of HepAD38 cells triggered 514 DEGs (**Fig. 2A**). Of the ICAA DEGs identified, 253 overlapped with the HBV DEGs. Of these, 189 were oppositely regulated by ICAA and HBV, while 64 were regulated in the same direction (**Fig. 2A**).

**Fig. 2.**
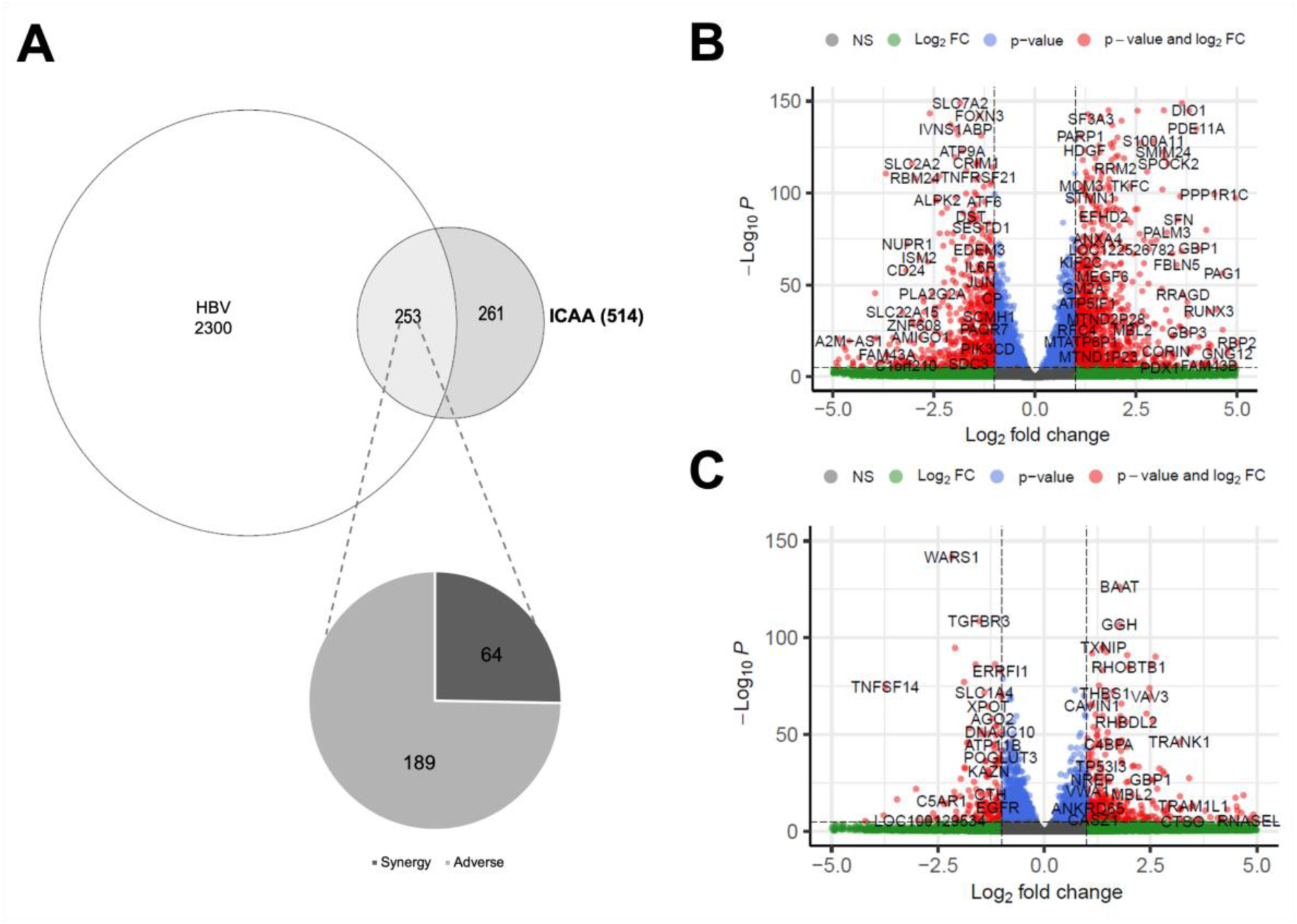
Transcriptomic analysis of HBV- and ICAA-associated differentially expressed genes (DEGs) in HepAD38 cells. (**A**) Gene counts of HBV- and ICAA-associated DEGs following normalization using DMSO-treated HepG2 cells as a reference. Differential expression analysis identified 2300 HBV-associated DEGs in DMSO-treated HepAD38 cells and 514 ICAA-associated DEGs in ICAA-treated HepAD38 cells at day 3 post-treatment. Among the 514 ICAA-associated DEGs, 253 overlapped with HBV-regulated genes, including 189 genes oppositely regulated by ICAA and HBV, whereas 64 genes were regulated in the same direction. The remaining 161 DEGs were specifically induced by ICAA. (**B–C**) Volcano plots showing significantly downregulated and upregulated DEGs (p < 0.05) associated with HBV expression (**B**) and ICAA treatment (**C**), respectively.

Moreover, Heme Oxygenase-1 (HO-1) was among the 64 DEGs of the same modulation pattern and was found to be strongly upregulated following ICAA treatment, consistent with our previously published data [14]. This finding indicates that HBV-expressing HepAD38 cells induce HO-1 as an antioxidant response to HBV, while ICAA treatment further enhances this response through additional HO-1 upregulation. The remaining 161 ICAA-associated DEGs were induced specifically by ICAA. Analysis of the 189 oppositely induced genes showed a strong downregulation pattern by HBV, while an upregulated reversal pattern was seen in ICAA-treated cells. General statistical analysis of the overall transcripts presented in **Table 2** shows a reversal effect.

**Table 2.** Summary statistics of differential gene expression 3 days post-treatment of HepAD38.

| Inducer | Min | 1st Qu. | Median | Mean | 3rd Qu. | Max |
| --- | --- | --- | --- | --- | --- | --- |
| HBV | -6.588 | -2.775 | -1.858 | -1.595 | -1.168 | 4.995 |
| ICAA | -3.979 | 1.070 | 1.487 | 1.413 | 2.119 | 5.417 |

Additionally, volcano plots were generated to visualize differentially expressed genes based on the magnitude of the expression change (log₂ fold change) and statistical significance (adjusted p-value). Those for HBV- and ICAA-regulated DEGs are shown in **Fig. 2B** and **2C**, respectively. The plots present significantly upregulated and downregulated genes. Overall, these data revealed clear differences in transcription between the study conditions, highlighting the importance of focusing downstream analysis primarily on DEGs from the three-day post-treatment period in HepAD38 cells.

### 2.2. Enriched pathways of ICAA’s DEGs are mainly associated with tumorigenesis

In addition to focusing our downstream analysis on HepAD38 cells 3 days post-treatment, we aimed to investigate the biological relevance of ICAA-associated DEGs in the context of HBV with respect to proliferation or cell death. To contextualize the identified DEGs, we performed KEGG-based enrichment analyses.

The ICAA-associated DEGs were classified into two categories: (i) oppositely regulated genes, which are defined as genes that exhibit inverse regulation patterns between the expression profiles of HBV and ICAA, and (ii) ICAA-specific genes, which are defined as genes that are differentially expressed exclusively following ICAA treatment in HepAD38 cells. The enriched pathways were largely related to viral carcinogenesis and cancer-related signaling pathways, including apoptosis, multiple species, MAPK, p53, and transcriptional misregulation in cancer (**Fig. 3A**).

**Fig. 3.**
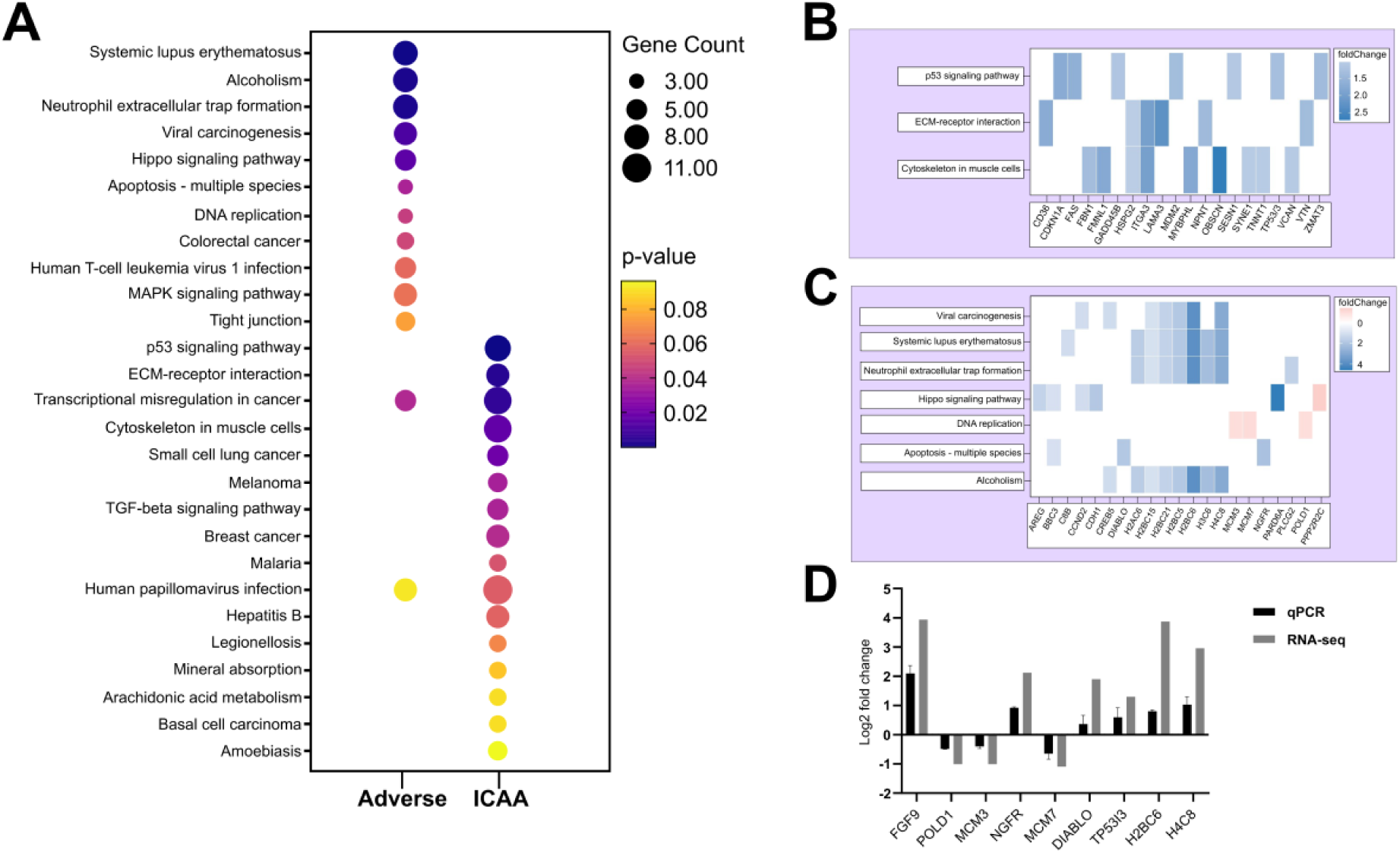
KEGG pathway enrichment analysis of oppositely regulated and ICAA-specific DEGs. (**A**) The graph illustrates significantly enriched KEGG pathways identified from oppositely regulated DEGs (“adverse”), representing genes inversely regulated between HBV-associated and ICAA-treated conditions in HepAD38; and ICAA-specific DEGs (“ICAA”), corresponding to genes uniquely modulated following ICAA treatment in HepAD38 cells. Each dot represents an enriched pathway, with color intensity corresponding to enrichment significance (*p*-value) and dot size proportional to the number of genes associated with each pathway. (**B**) and (**C**) are gene sets extracted from significantly enriched KEGG pathways identified from ICAA-specific and oppositely regulated DEGs, respectively in HepAD38. Shared genes across multiple enriched pathways are displayed to illustrate common molecular regulators contributing to HBV-associated transcriptional programs reversed following ICAA treatment or modulation by ICAA independently of HBV-associated regulation. (**D**) Validation of selected identified RNA-seq DEGs by qPCR. Relative gene expression levels obtained by qPCR were first calculated as fold-change relative to the control and were then transformed to log2 values for comparison with transcriptomic data. The qPCR analysis confirmed the expression trends observed in the RNA-seq dataset, supporting the reliability and reproducibility of the identified DEGs.

To identify key genes within the differentially expressed genes (DEGs) of the enriched pathways that may function as regulators of the biological response and provide insight into the mechanisms underlying ICAA- and HBV-associated cellular changes, a hub gene analysis was performed. Analysis of ICAA-specific transcripts were mainly involved in cell cycle regulation (**Fig. 3B**), suggesting an additional HBV-independent proliferative response. Whereas the oppositely regulated DEGs revealed that it was predominantly enriched in histone-related genes, indicating a reversal of HBV-associated epigenetic modulation (**Fig. 3C**). These results suggest that ICAA modulates both HBV-associated chromatin organization and broader cell cycle control mechanisms. Furthermore, the qPCR-based expression profiles of nine randomly selected genes (six of which were upregulated and three of which were downregulated) correlated with the RNA-seq data, confirming the reliability of the generated RNA-seq data (**Fig. 3D**).

### 2.3. Identified pathways converge to cell cycle regulation

As previously described, ICAA treatment of HepAD38 led to oppositely induced DEGs and ICAA-specific DEGs involving epigenetic and cell cycle modulations, respectively. To obtain a more meaningful overview of the identified transcripts in a biological context, we then used the Enrichr online tool (https://maayanlab.cloud/Enrichr/) to generate a Uniform Manifold Approximation and Projection (UMAP) representation of the KEGG-enriched pathways. Since a UMAP generates a neighborhood graph of the enriched pathways, we generated Venn diagrams to visualize the overlap between the selected clusters of pathways. For the oppositely regulated genes (**Fig. 4A**), we selected two clusters of pathways for overlap analysis: one involving colorectal cancer, MAPK and Hippo signaling, and the other including viral carcinogenesis, alcoholism, neutrophil extracellular trap formation, and transcriptional misregulation in cancer. The overlapping genes of these clusters were further investigated for their biological relevance. The same analysis was extended to ICAA-specific genes, using two clusters: one containing p53 signaling, ECM-receptor interactions, melanoma, and breast cancer, and the second including legionellosis and malaria (**Fig. 4B**). In both cases, the overlapping genes were primarily associated with chromatin organization (H2AC6, H2BC5/6/15/21, H4C8, H3C6), cell cycle regulation and proliferation (CCND2, CDKN1A, FOS, AREG, EREG), and stress/apoptotic signaling (BBC3, GADD45B, DDB2, CXCL8, TLR4).

**Fig. 4.**
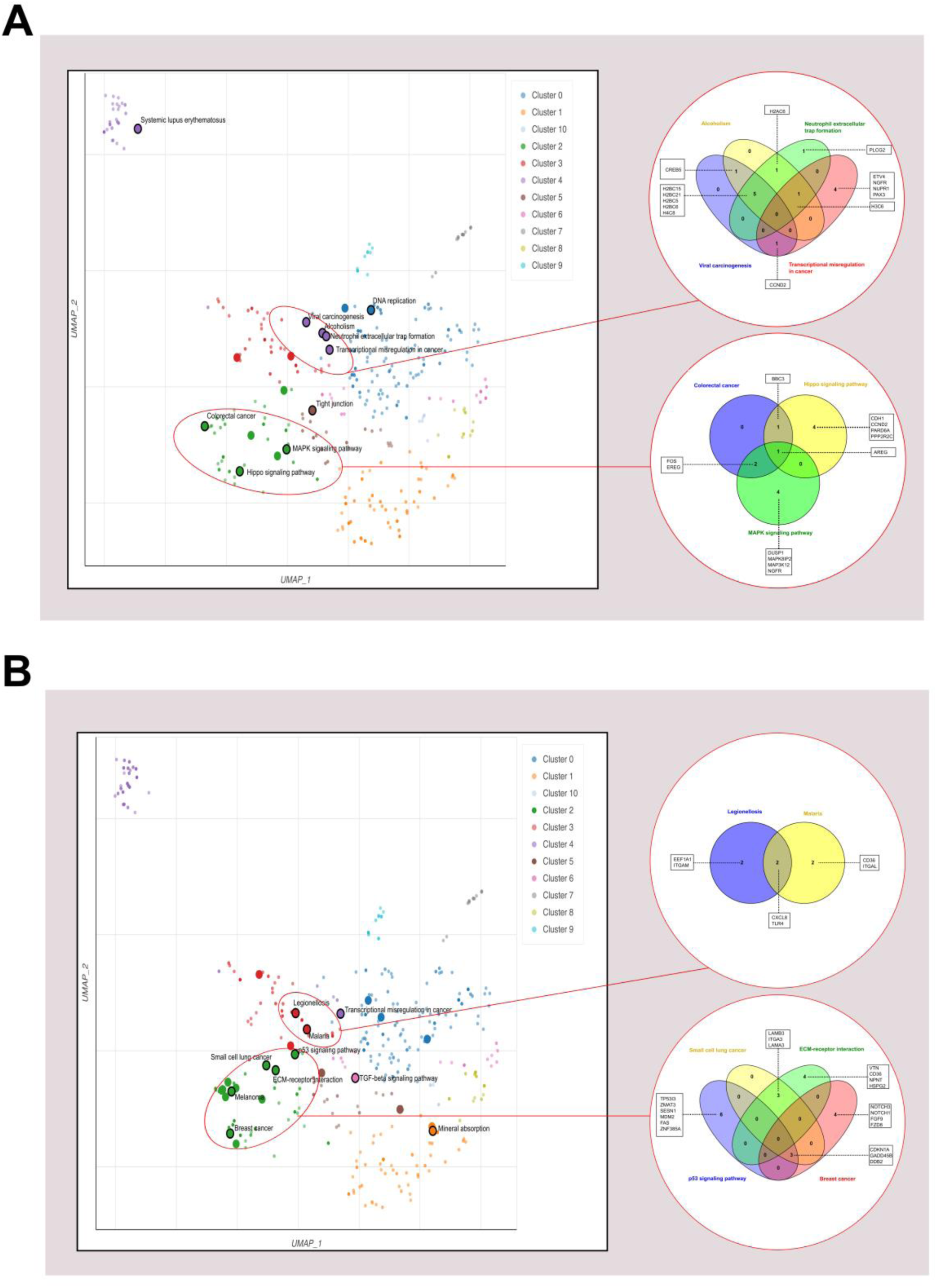
UMAP visualizations of KEGG-enriched pathways generated by Enrichr. (**A**) Enriched pathways associated with oppositely regulated genes. The pathways were clustered according to their shared genes, and the selected clusters were further integrated into Venn diagrams. Two main clusters of pathways have been identified: (i) viral carcinogenesis, alcoholism, neutrophil extracellular trap formation, and transcriptional misregulation in cancer pathways, and (ii) colorectal cancer, MAPK, and Hippo signaling pathways; **(B)** Enriched pathways associated with ICAA-specific DEGs. Pathways were clustered according to common genes, and the selected clusters were further integrated into Venn diagrams. Two main clusters of pathways have been identified: (i) p53 signaling, ECM-receptor interaction, breast cancer, and small cell lung cancer; and (ii) legionellosis and malaria.

### 2.4. Identified transcription factors are associated with apoptosis, oxidative stress, DNA repair, and cellular adaptation to stress

Although pathway enrichment followed by hub gene analysis identified biological processes (epigenetics, cell cycle, and stress/apoptotic signaling) in ICAA-treated HepAD38 DEGs, the large number of altered genes (514 ICAA-induced DEGs) makes mechanistic interpretation challenging. Therefore, we performed transcriptional regulator (TR) analysis to identify key upstream regulators that could account for the observed ICAA-induced DEGs in HepAD38 cells. This narrowed the complexity of the dataset and provided insight into the DEGs. We employed the TRRUST (Transcriptional Regulatory Relationships Unraveled by Sentence-based Text mining) database in the Metascape online tool (https://metascape.org). The results identified transcription factors including TP53, TP73, TP63, FOXO3, and ATF4, which suggest cellular stress and DNA damage responses (**Supplemental Fig. 1**). These regulators are generally known to regulate transcriptional programs involved in apoptosis, oxidative stress, DNA repair, and cellular adaptation to stress. TP53 family members are particularly central mediators of genotoxic stress signaling. FOXO3 and ATF4 are associated with oxidative stress and integrative stress response pathways [25–27]. These data indicate an activation of stress-related regulatory networks, which may reflect an increase in cellular stress and DNA damage-associated signaling.

### 2.5. Overlay of differentially expressed genes highlights a strong p53 pathway modulation

We decided to investigate p53-associated pathways due to the identification of TP53 family members, as along with FOXO3 and ATF4, which are key regulators of cell cycle arrest, apoptosis, and DNA damage. We analyzed the p53 pathway using DEGs for all three conditions: DMSO-treated HepAD38 cells, ICAA-treated HepAD38 cells, and ICAA-treated HepG2 cells three days post-treatment. For this, the p53 pathway was fed with data consisting of (i) DEGs associated with HBV in HepAD38 cells (HBV effects), (ii) DEGs induced by ICAA in HepAD38 cells (ICAA effects in the presence of HBV expression), and (iii) DEGs induced by ICAA in HepG2 cells (ICAA effects in the absence of HBV expression). Overlaying the gene expression data of the aforementioned datasets onto the KEGG p53 signaling pathway using color-coded nodes allows visualization of the differences in the regulation pattern of the components of the pathway (**Fig. 5A**). The Pathview result shows that HBV downregulates p53 gene expression, while ICAA treatment normalizes it. Western blot analysis confirmed an increased amount of p53 protein upon ICAA treatment in HepAD38 cells (**Fig. 5B**). Furthermore, several upstream and downstream components of the pathway were oppositely regulated between the HBV and ICAA conditions, including (MDM2, MDM-X, CHK1, Gadd45, Cyclin D/CDK4/6, Cycline E, PUMA, Bcl2, KAI, and Sestrins). Given the strong dysregulation of p53 downstream pathways and the centrality of p53 as a key regulator of cell cycle arrest, apoptosis and DNA damage, we hypothesize that the observed changes are associated with modulation of apoptosis in HepAD38 cells.

**Fig. 5.**
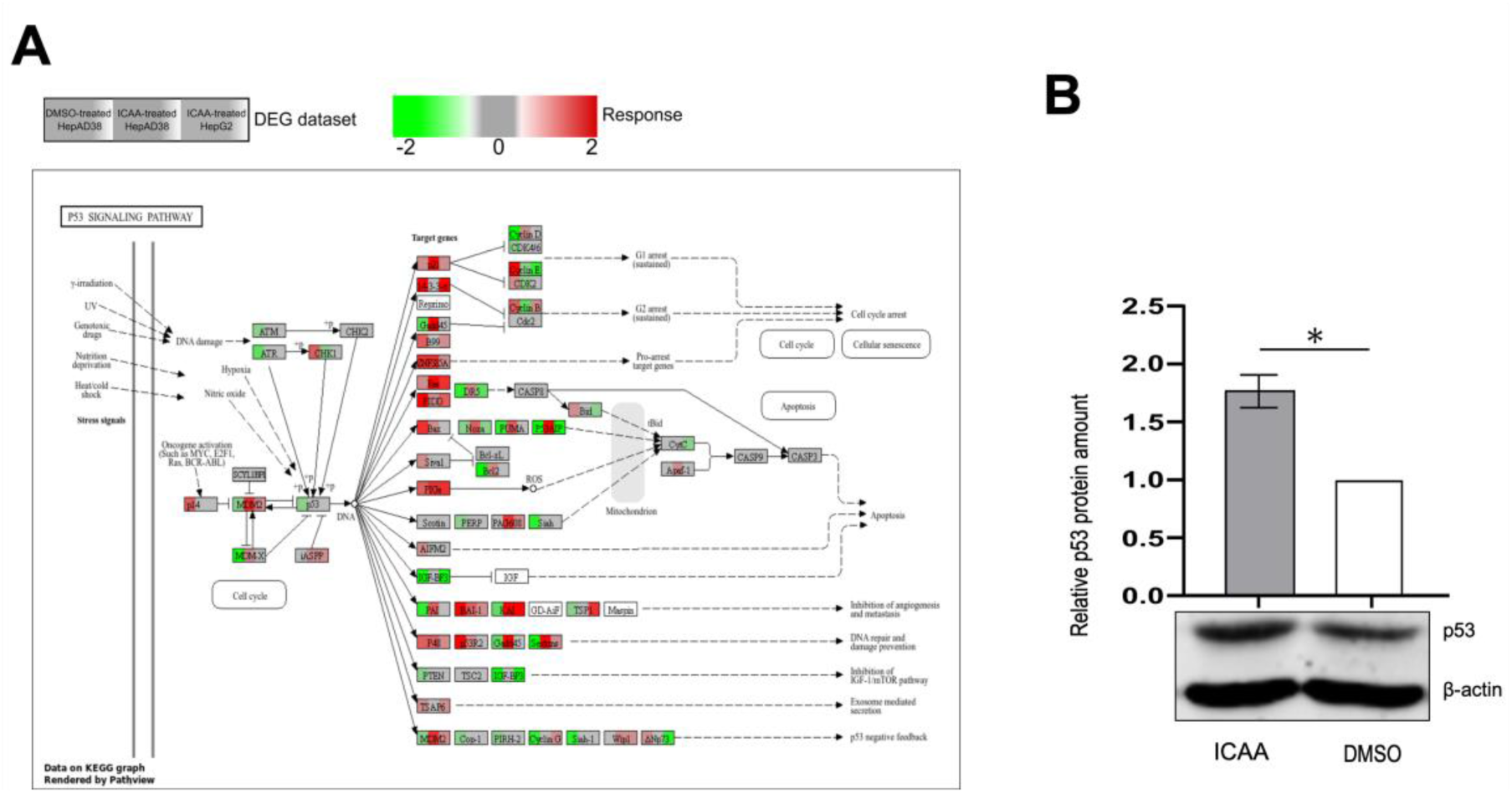
**Differentially expressed genes in the p53 pathway**. (**A**) KEGG Pathview of the p53 signaling pathway. The Pathview illustrates the central role of p53 as a tumor suppressor, which regulates cell cycle arrest, DNA repair, senescence and apoptosis in response to cellular stress. Each gene of the pathway was presented in three blocks as log2FC of (i) DEGs associated with HBV in HepAD38 (HBV effect), (ii) DEGs induced by ICAA in HepAD38 (ICAA plus HBV), and (iii) DEGs induced by ICAA in HepG2 (ICAA without HBV), respectively. The log2FCs were represented by a color gradient to indicate upregulated and downregulated genes. (**B**) Representative image for the Western blot showing the p53 protein amount 3 days post-treatment in HepAD38 and the quantification. The p53 signal intensity was normalized to actin. Three independent biological replicates (n = 3) were used for statistical analysis. Data sets were analyzed using an unpaired, two-tailed Student’s t-test: \**p* < 0.05.

### 2.6. ICAA promotes the activation of downstream apoptotic markers

We observed various DEGs (MDM2, MDM-X, CHK1, Gadd45, Cyclin D/CDK4/6, Cycline E, PUMA, Bcl2, KAI, and Sestrins) that are associated with pathways controlling apoptosis. There was also a strong dysregulation of p53 downstream pathways, which are key regulators of cell cycle arrest, apoptosis, and DNA damage repair. Previous studies have described that HBV primarily induces survival to evade cell death by promoting proliferative pathways [28]. Based on this, we investigated apoptosis markers in HepAD38 cells. Apoptosis analysis using annexin V staining identified a subset of apoptotic cells in both ICAA-treated and-untreated cells (**Fig. 6A**). For a more detailed analysis, western blot was performed to investigate caspase-9- and PARP-cleavage. The western blot analysis of untreated HepAD38 cells revealed cleaved caspase-9, but no significant activation of PARP, suggesting induction of the intrinsic apoptotic pathway with concomitant blockade of apoptotic execution (**Fig. 6B**). In contrast to this, ICAA-treatment of HepAD38 cells inhibited caspase-9 cleavage in a dose-dependent manner while activating the executive apoptotic enzyme, Poly(ADP-ribose) polymerase (PARP) (**Fig. 6B**). Furthermore, apoptotic DNA fragmentation was predominantly identified in ICAA-treated HepAD38 cells (**Fig. 6C**). These results suggest that HBV primarily activates the intrinsic apoptosis pathway, more likely by affecting mitochondrial function, which leads to an elevated ROS level. However, it blocks the subsequent execution to avoid cell death. In contrast, ICAA-treated cells showed reduced caspase-9 activation, indicating a dose-dependent antioxidant response combined with cleaved PARP, which suggests a rescued extrinsic apoptosis pathway. Taken together, these findings imply that HBV alters cellular regulatory pathways, including the cell cycle and apoptotic signaling, to maintain cell survival and effectively evade cell death. ICAA treatment appears to restore these cellular functions, potentially restoring effective cell death mechanisms modulated by HBV.

**Fig. 6.**
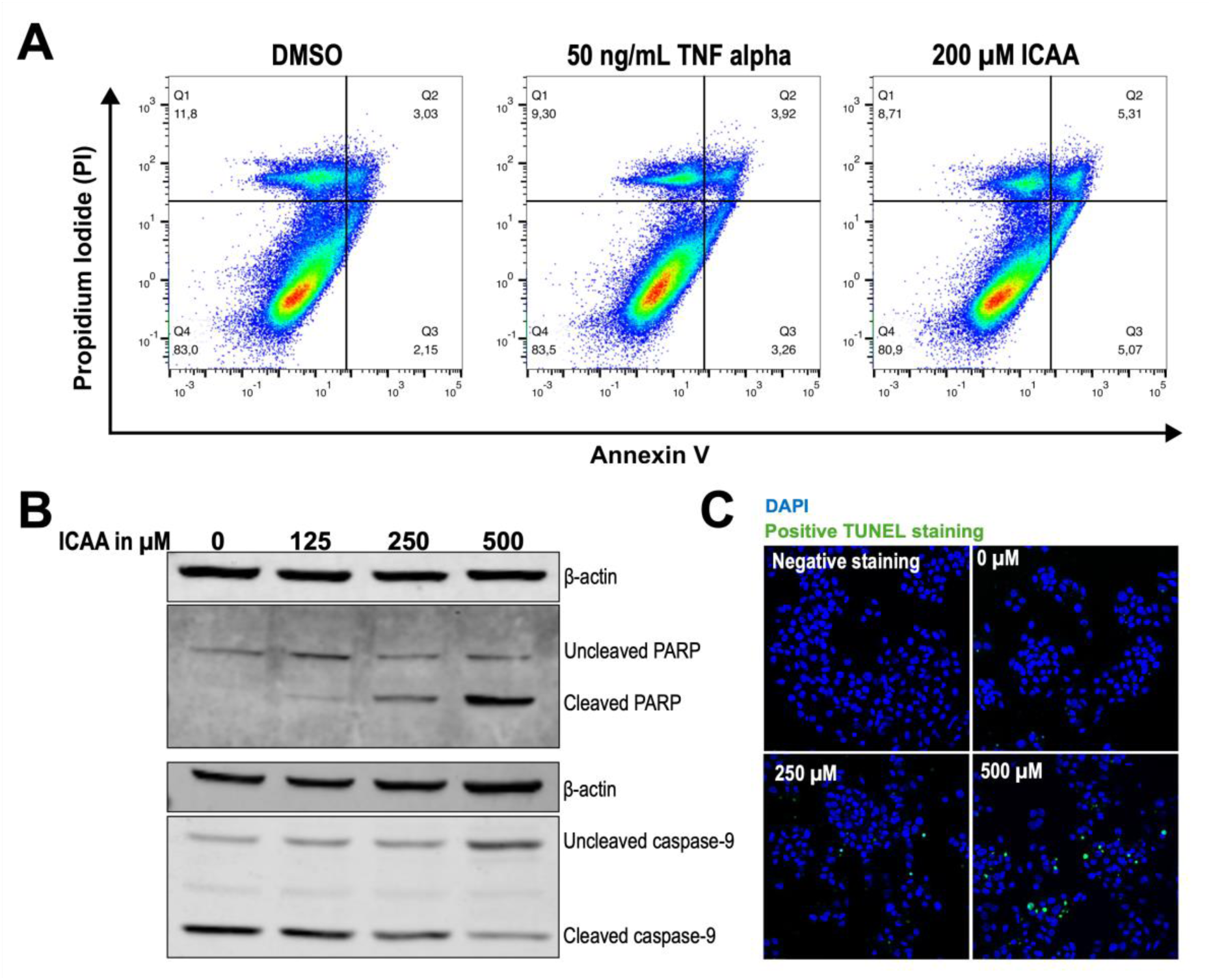
**Investigation of apoptotic markers in HepAD38**. (**A**) Annexin V-stained cells were analyzed for non-apoptotic cells (Q4), early and late apoptosis (Q3) and (Q2), respectively, and necrotic cells (Q1). TNF-alpha at a concentration of 50 ng/mL was used as control. (**B**) Western blot showing a dose-dependent cleavage of caspase-9 and PARP in HepAD38 3 days post-treatment with 0-500 µM ICAA. (**C**) TUNEL assay for staining apoptotic DNA fragmentation; nuclei were stained in blue using DAPI, and the green channel shows apoptotic DNA fragmentation.

Furthermore, a TUNEL assay identified apoptotic DNA fragmentation in a dose-dependent manner, providing proof of apoptosis in ICAA-treated HepAD38 cells.

### 2.7. ICAA-induced cell death involves a non-canonical apoptotic mechanism

To further investigate whether the observed ICAA-induced rescue of apoptosis involves a canonical apoptotic mechanism, we analyzed the DEGs from ICAA-treated HepG2 cells three days post-treatment for apoptosis-related genes. However, KEGG pathway enrichment analysis for DEGs identified in the ICAA-treated HepG2 cells revealed no direct modulation of canonical apoptosis-related pathways (**Supplemental Fig. 2**). However, the enriched pathways indicated a strong modulation of hepatic metabolism and detoxification processes. These processes include cytochrome P450 xenobiotic metabolism, drug metabolism, hepatic oxidative stress, retinol metabolism, bile acid biosynthesis, cholesterol metabolism, and PPAR signaling (**See data availability section, for full list**). This reflects an impact on hepatic metabolic function and oxidative stress control.

Based on this observation, we reanalyzed the enriched pathways in HepAD38 cells. Of the 2300 DEGs from DMSO-treated HepAD38 cells, apoptosis pathways were not among the top-ranked pathways. Nevertheless, they were present in the full enrichment list (**See data availability section**) and comprised 28 genes (**Supplemental Fig. 3**). When we analyzed the 514 DEGs from ICAA-treated HepAD38 cells, however we identified 8 apoptosis-related genes were identified, including BBC3 (PUMA), CTSS, DIABLO, FAS, FOS, GADD45B, PRF1, and TUBA1A. Comparison with the 28 apoptosis-associated genes identified in DMSO-treated HepAD38 cells revealed no direct overlap at the individual gene level. Although pathway visualization suggested common CTS and TUBA nodes, these corresponded to different family members in each dataset (CTSS versus CTSO and TUBA1B /TUBA1C/TUBAL3, respectively). Thus, the apparent overlap reflects shared gene families rather than direct oppositely regulated gene expression (**Supplemental Fig. 3**). None of these genes were induced in ICAA-treated HepG2 cells.

Overall, these data suggest that ICAA-induced cell death in HBV-expressing cells occurs through a non-canonical apoptotic mechanism. ICAA selectively affects the metabolic and survival adaptations of HBV, restoring apoptotic capability and leading to execution of cell death in HBV-expressing HepAD38 cells.

### 2.8. Modulation of ER protein processing pathways

ICAA has been shown to reverse and regulate several transcripts induced by HBV in HepAD38 cells. This includes the dysregulation of apoptotic pathways, which may contribute to HBV persistence and replication. ICAA has previously been shown to interfere with several steps of the HBV life cycle, including viral maturation. We investigated transcriptomic changes related to protein processing, a process which involves disulfide bond formation and folding control in the endoplasmic reticulum (ER). Both of these processes are required for HBV maturation. Using the data set as described in **Fig. 5** and the Pathview module in KEGG, we found an increased transcriptional expression of genes involved in translocation, folding, trafficking, and suppression of the ER stress response by HBV. This reflects the formation of viral proteins and virus assembly (**Fig. 7**). However, ICAA treatment reversed a subset of HBV-induced changes. This suggests that ER functions required for the effective replication and maturation of HBV may be impaired by ICAA treatment. Key oppositely induced genes included: secretory pathway proteins 61/62/63 (Sec61/62/63) and Glutathione S-transferases (GSTs) which contribute to proper protein folding and prevention of aggregation, Glucosidase I/II (GlcI/II) (involved in protein folding), ATF6 (an unfolded protein response (UPR)/ER-stress regulator), Sec23/24 (involved in ER-to-Golgi transport,), and heat shock proteins 40/70 (Hsp40/70) which are important for protein folding and degradation in the ER [29–34]. These data support previously described effects of ICAA on HBV, such as affecting disulfide bond formation and impairing viral maturation. This suggests that reactivation of executionary apoptosis in HepAD38 cells following ICAA treatment may not be a direct effect, but rather the consequence of impaired viral maturation and restored cellular functions.

**Fig. 7.**
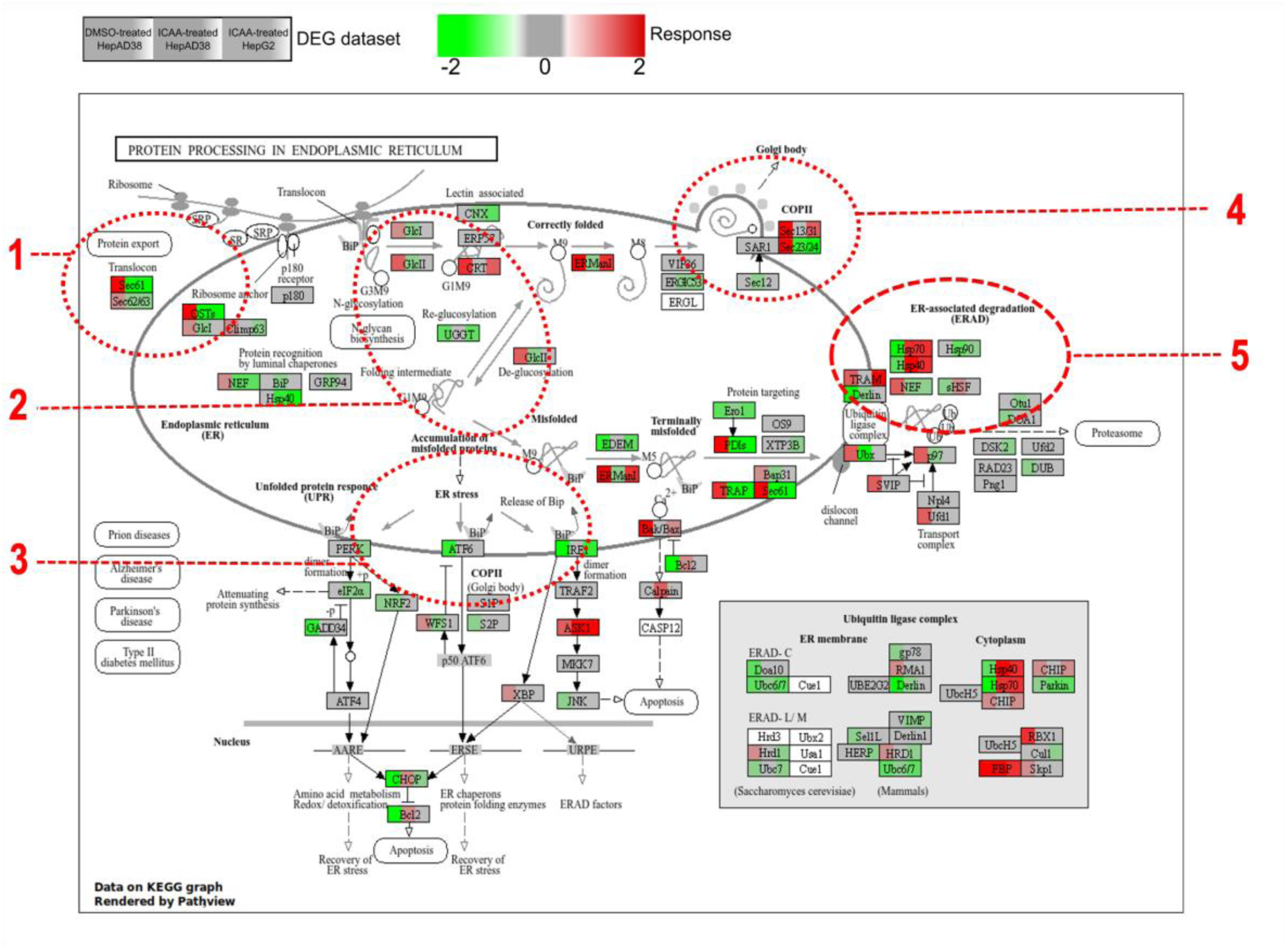
**KEGG Pathview of protein processing in the endoplasmic reticulum (ER)**. Each gene of the pathway was presented in three blocks as log2fc of (i) DEGs associated with HBV in HepAD38 (HBV effect), (ii) DEGs induced by ICAA in HepAD38 (ICAA plus HBV), and (iii) DEGs induced by ICAA in HepG2 (ICAA without HBV), respectively. The log2FC were represented by a color gradient to indicate upregulated and downregulated genes. The Pathview shows the molecular machinery responsible for folding, maturation, quality control, and trafficking of proteins in the ER. This invokes the following: (**1**) The translocation of proteins to the endoplasmic reticulum allows newly synthesized proteins to enter the lumen of the ER for further processing and maturation. (**2**) In the ER, proteins are folded and glycosylated to achieve the correct structure and function. (**3**) The accumulation of unfolded or misfolded proteins activates ER stress signals and the unfolded protein responses (UPR) in order to restore ER homeostasis. (**4**) Well-formed proteins are packed in COPII-coated vesicles and transported from the ER to the Golgi apparatus where they mature and are secreted. (**5**) Misfolded proteins are eliminated to maintain protein quality control.

### 2.9. Functional annotation analysis links identified DEGs to apoptosis-inducing drugs

A functional annotation analysis was conducted using the DAVID bioinformatic tool to investigate the biological relevance of the 514 ICAA DEGs in the functional context of known drugs. The analysis revealed that the identified DEGs correlate with the signatures of substances known to affect apoptosis, such as Lestaurtinib, Kw-2449, Fedratinib, Sunitinib and Staurosporine (**Table 3**). These results further support the hypothesis that the executory apoptosis process was rescued upon ICAA treatment in HBV-expressing cells.

**Table 3.**
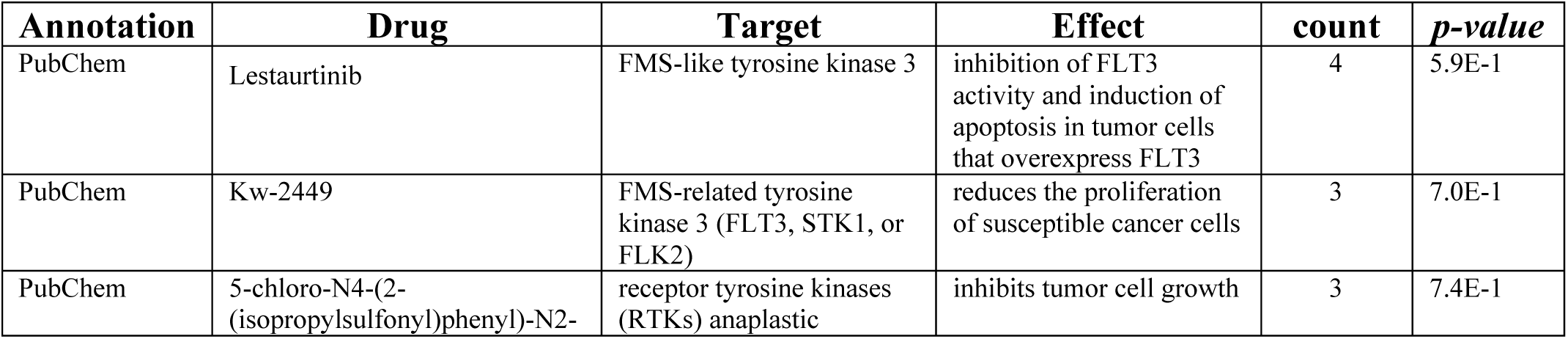
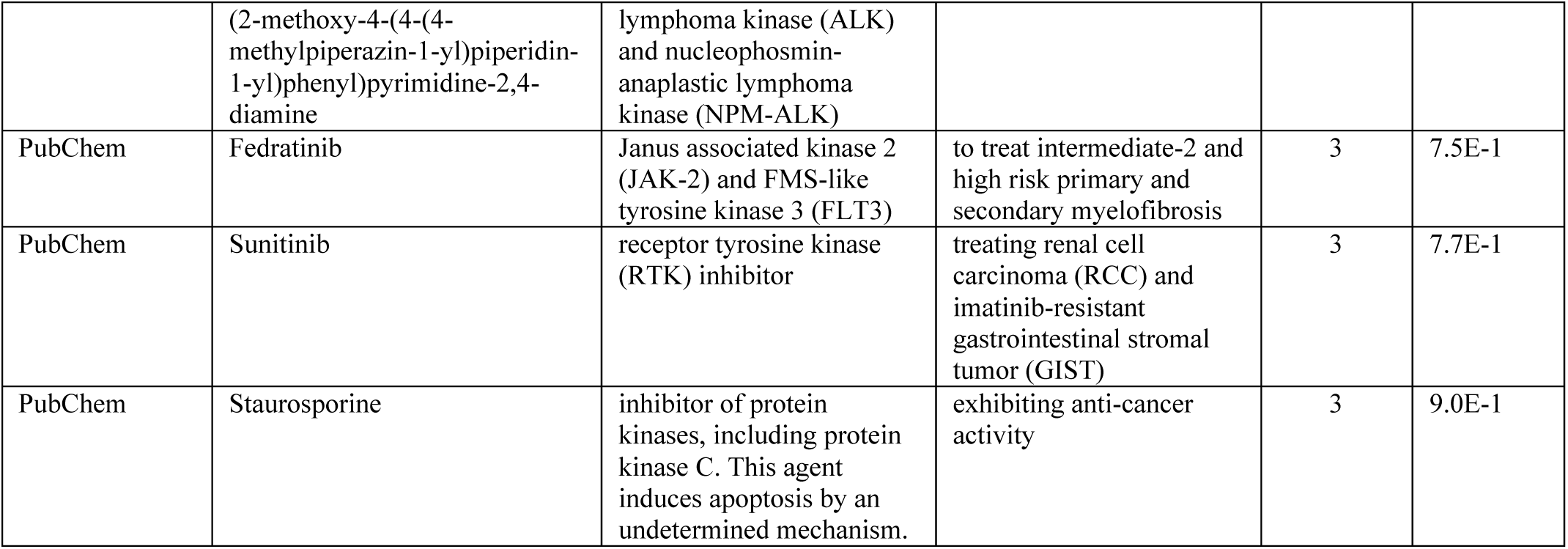
Key drugs identified in PubChem through function annotation in DAVID bioinformatic.

## 3. Discussion

Hepatocarcinogenesis associated with HBV is a multi-factorial process driven by chronic inflammation, persistent viral replication, oxidative stress, and progressive dysregulation of the host transcriptional program [35–37]. This study’s transcriptomic profiling suggests extensive HBV-mediated modulation of host gene expression in HepAD38 cells, with approximately 2300 DEGs as compared to DMSO-treated HepG2 cells. In contrast, ICAA induced a more selective transcriptional response involving 514 DEGs in HepAD38 cells as compared to DMSO-treated HepG2 cells. The significantly wider transcriptional impact of HBV in HepAD38 cells (using HepG2 cells as a reference) is consistent with previous reports showing that HBV infection globally alters the transcriptional and epigenetic landscape of the host, creating a favorable environment for virus persistence and carcinogenesis. PCA on the generated DEGs showed clear time-dependent transcriptional changes after ICAA treatment, with the clearest effect observed at 3 days post-treatment in HepAD38. This finding suggests that the molecular effects of ICAA are gradual and may take a sufficient period of time to overcome the transcriptional control mechanisms mediated by HBV. Of the 514 identified ICAA DEGs in HepAD38, 253 overlapped with HBV-associated genes. Importantly, most of the overlapping genes (189) were oppositely regulated by HBV and ICAA, while 64 genes were similarly regulated but to a different magnitude. These observations strongly suggest that ICAA reverses a subset of transcriptional changes induced by HBV, rather than simply inducing a non-specific cellular response. Among the 64 DEGs similarly induced by ICAA and HBV, included HO-1, which was strongly upregulated following ICAA treatment. This is consistent with our previously published data [14], indicating that HBV-expressing HepAD38 cells induce HO-1 as an antioxidant response to HBV, while ICAA treatment further enhances this response through additional HO-1 upregulation. In addition, the identification of ICAA-specific DEGs (261) in HepAD38 cells suggests that the compound also activates independent transcriptional programs in addition to directly reversing HBV-induced effects.

Functional enrichment analysis further supports the biological relevance of these findings. KEGG pathway analyses of HepAD38 cells show enrichment of pathways related to viral carcinogenesis, apoptosis, MAPK signaling, p53 signaling, and transcriptional misregulation in cancer. These pathways are of high relevance for the HBV life cycle and HBV-associated pathogenesis, including HCC formation [35]. HBV, particularly HBx, has been shown to activate the MAPK and PI3K-AKT signaling pathways [28], which promote survival, proliferation, and resistance to apoptosis in hepatocytes. Persistent deregulation of these pathways contributes to chronic hepatic impairment and malignant transformation, i.e. by a tumor promoter-like function as described for the HBV PreS2 activator MHBs^t^ [38].Therefore, identification of MAPK and cancer-related pathways in the oppositely regulated genes suggests that ICAA may disrupt the signaling networks that are relevant for HBV-induced oncogenesis. Interestingly, the analysis of the hub genes in HepAD38 revealed that oppositely regulated DEGs were enriched mainly in histone-related pathways. Epigenetic regulation has also been identified as a key mechanism of HBV persistence and hepatocellular carcinogenesis [39, 40]. In addition, HBV replication relays on cccDNA’s transcriptional activity, which is tightly regulated by histone modification and chromatin remodeling [40]. HBx has been shown to modify histone acetylation and methylation pathways in order to maintain viral transcription and suppress the host antiviral response [41]. Thus, the enrichment of HBV-dependently modulated epigenetic plus its opposite regulation by ICAA, suggests a regulatory effect of ICAA on disrupted epigenetics during HBV replication. Moreover, Hippo signaling pathway was also enriched. The Hippo pathway is increasingly recognized as a major regulator of hepatic volume, tissue regeneration, and the development of hepatocellular carcinoma [42, 43]. Dysregulation of Hippo signaling promotes uncontrolled proliferation of hepatocytes and resistance to apoptosis, processes that are commonly observed in hepatic disease associated with HBV[44, 45]. Enrichment of this pathway thus supports the hypothesis that ICAA modulates key oncogenic pathways activated by chronic HBV infection.

ICAA-specific transcripts were enriched mainly in pathways related to p53 signaling and regulation of the cell cycle. The p53 pathway is of particular importance in HBV-induced carcinogenesis, since p53 is a central regulator of genomic stability, cell cycle arrest, senescence, and apoptosis. Thus, enrichment of p53 signaling together with several oppositely regulated genes (MDM2, MDM-X, CHK1, Gadd45, Cyclin D/CDK4/6, Cycline E, PUMA, Bcl2, KAI, Sestrins.) suggests that ICAA promotes tumor suppressor function impaired by HBV.

One of the most important findings of this study concerns apoptosis regulation. HBV has been described to have paradoxical effects on apoptotic signaling pathways [46]. While HBV infection can induce oxidative stress, mitochondrial dysfunction, and DNA damage [47, 48] that may activate apoptosis, the virus simultaneously suppresses the downstream pathways of apoptosis, thereby conserving the survival of infected hepatocytes. This equilibrium allows for continued viral replication while avoiding premature elimination of infected cells. Several studies have shown that HBx can activate apoptotic signals by inducing mitochondrial ROS and releasing cytochrome c, while simultaneously inhibiting the activation of downstream caspases by modulation of transcription factor NF-κB, vesicle-derived kinase, and anti-apoptotic BCL-2 [47, 48]. Apoptosis data obtained in this study support this model. Annexin V staining revealed apoptotic populations in both untreated and treated HepAD38 cells, suggesting that HBV replication itself induces apoptotic stress. However, western blot analyses revealed markedly different apoptotic profiles in the untreated and treated HBV-expressing cells. In untreated HBV-expressing cells, extensive cleavage of caspase-9 was observed, while PARP was largely non-cleaved. Activation of caspase-9 is typical of an intrinsic mitochondrial apoptotic pathway and is consistent with previous reports showing mitochondrial dysfunction and ROS accumulation induced by HBV [47, 48]. However, the absence of downstream PARP cleavage suggests that HBV might block the terminal apoptotic pathway, despite activation of the downstream apoptotic signals. This incomplete apoptotic phenotype may be a viral strategy for maintaining cell survival under chronic stress conditions. In contrast, treatment with ICAA reduced cleavage of caspase-9, while inducing dose-dependent cleavage of PARP. These findings indicate that ICAA may exert an antioxidant effect and favor the apoptotic execution process [49]. A positive TUNEL assay further confirmed effective cell death induction. The apoptosis observed following ICAA treatment appears paradoxical given the strong antioxidant response of ICAA. Antioxidants are generally associated with reduced oxidative stress and suppression of ROS-mediated apoptosis signaling [50–52]. Although ICAA exhibited antioxidant effects, the observed induction of apoptosis is not necessarily contradictory. The apoptotic rescue effect may not result from direct canonical cell death pathway but rather from restoration of cellular homeostasis in HBV-replicating HepAD38. One possible explanation could be the inhibition of HBV replication and/or maturation by ICAA, thereby relieving the HBV-mediated suppression of cellular regulatory mechanisms. As a result, HepAD38 may regain its ability to undergo physiological apoptosis to facilitate the elimination of HBV-positive cells. This can be correlated with the observation that p53 expression is downregulated in HepAD38, while a restored p53 expression level in ICAA-treated HepAD38 is comparable to that of HepG2. Therefore, the observed apoptotic rescue may reflect normalization of cellular quality control processes rather than a direct cytotoxicity. Together, these results indicate that both antiviral activity and restoration of cellular homeostasis may contribute to the transcriptional response induced by ICAA.

The use of a stable HBV expressing cell line instead of HBV-infected cells might appear as a limitation of our study. We decided for several reasons to use this system: In all infection-based models, only a portion of the cells are infected, which makes it difficult to identify differentially regulated genes due to the background of uninfected cells. When using PHHs, there is also a high degree of variability among the different isolates. Furthermore, the aim of the studies was to mimic the effect of ICAA on a stable, established infection (chronic HBV infection). A stable cell line comes much closer to this than an infected cell culture that expresses HBV only for a very limited period of time. Overall, these findings support a model where HBV induces a pro-survival state of the cell characterized by sustained downstream apoptotic stress, dysregulated control of the cell cycle, and suppression of the terminal apoptotic process. ICAA seems to counteract HBV-induced effects by reversing a subset of transcriptional changes, involving p53 and the cell cycle, which correlate with downstream apoptotic activation of apoptosis.

## 4. Conclusion

ICAA selectively reverses the metabolic and survival adaptations of HBV and restores apoptotic capability, leading to cell death in HBV-expressing HepAD38 cells.

These findings provide important mechanistic insights into how modulation of apoptosis and cell cycle regulation contributes to the survival of HBV-expressing cells. They also indicate that ICAA may have therapeutic potential in treating hepatic diseases and hepatocellular carcinogenesis associated with HBV.

## CRediT authorship contribution statement

**Giscard Wilfried Koyaweda**: Conceptualization, Methodology, Validation, Formal analysis, Investigation, Data Curation, Visualization, Writing - Original Draft, Writing - Review & Editing

**Mirco Glitscher**: Methodology, Investigation, Writing - Review & Editing

**Csaba Miskey**: Methodology, Validation, Data Curation, Visualization, Writing - Review & Editing

**Eberhard Hildt**: Conceptualization, Methodology, Resources, Supervision, Project administration, Funding acquisition, Writing - Original Draft, Writing - Review & Editing

## Acknowledgment

The authors thank the German Academic Exchange Service (DAAD) for providing scholarships to Giscard Wilfried Koyaweda during his PhD studies.

## Data availability statement

The raw sequencing files of the RNA-seq data generated in this study have been submitted to the NCBI SRA database (accession number PRJNAXXXXXXXXXXX). The accession number will be released upon acceptance of the manuscript and made publicly available.

The enriched pathways derived from the KEGG analysis used to generate the results presented in this study are available through the Mendeley Data repository (accession link: https://data.mendeley.com/preview/fv3dmnd9dc?a=759aa1e5-e68c-4288-922f-a817d7e0b423). These data will be made publicly available once the manuscript is accepted, at which point the repository link will be updated.

All other data supporting the findings of this study are available from the corresponding author upon request.

## Abbreviations

ICAA: Isochlorogenic acid A
HBV: hepatitis B virus
DMSO: dimethyl sulfoxide
DEGs: differentially expressed genes
KEGGP: Kyoto Encyclopedia of Genes and Genomes.
cccDNA: covalently closed circular DNA
cDNA: complementary DNA
DEGs: differentially expressed genes
DNA: deoxyribonucleic acid
DMSO: dimethyl sulfoxide
HBV: Hepatitis B virus
HCC: hepatocellular carcinoma
HO-1: heme oxygenase-1
ICAA: Isochlorogenic acid A
IFNs: interferons
KEGG: Kyoto Encyclopedia of Genes and Genomes
log2FC: log2 fold-change
NAs: nucleos(t)ide analogues
PCA: principal component analysis
PARP: poly(ADP-ribose) polymerase
qPCR: quantitative polymerase chain reaction
rcDNA: relaxed circular DNA
RNA: ribonucleic acid
RNA-seq: RNA sequencing
TNF alpha/TNF-α: tumor necrosis factor alpha

## Supplemental materials

**Supplemental figure 1.**
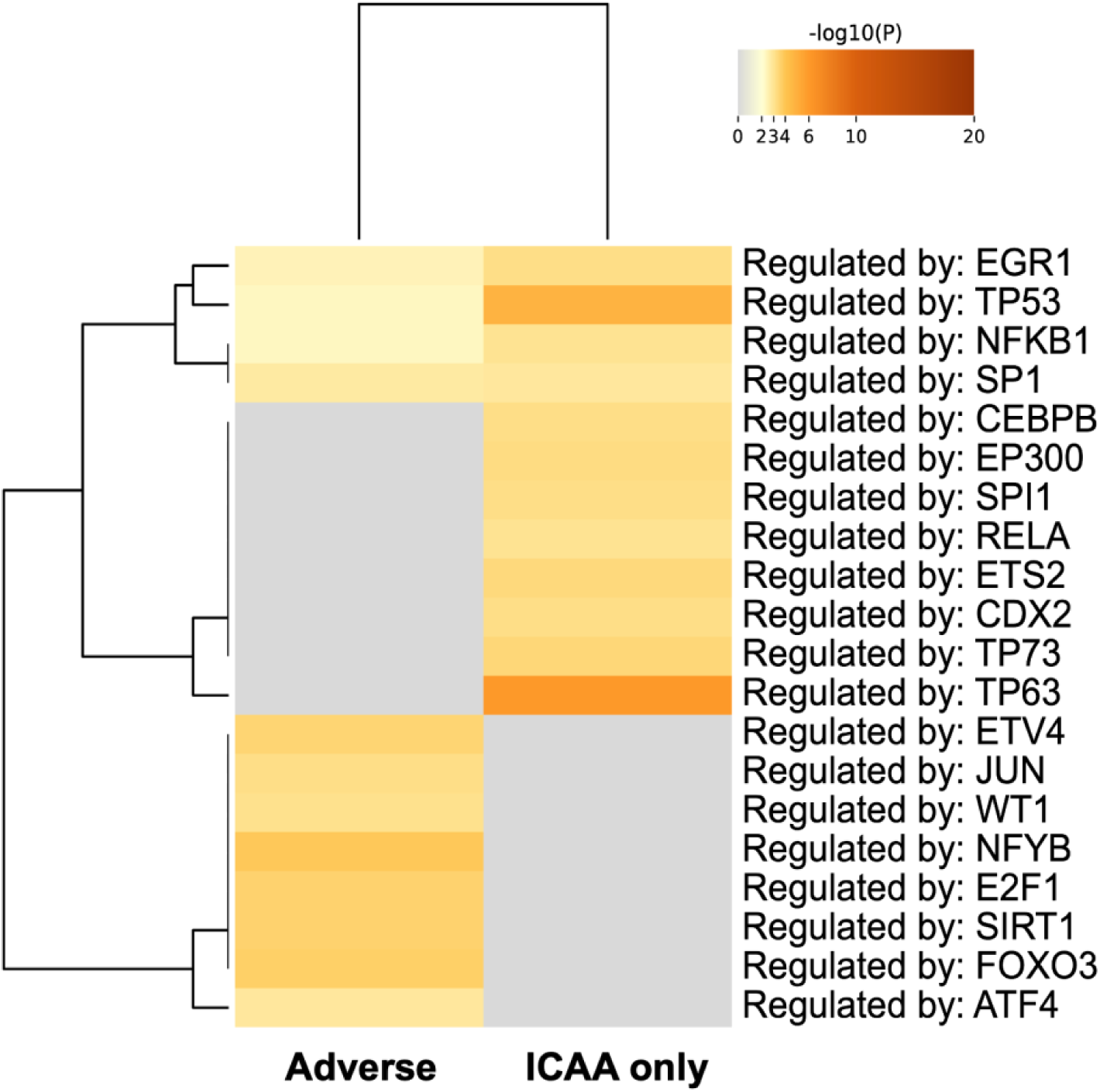
Enrichment of transcription factors. Transcription factor enrichment analysis based on the 514 ICAA-associated differentially expressed genes (DEGs) identified in HepAD38 cells.

**Supplemental figure 2.**
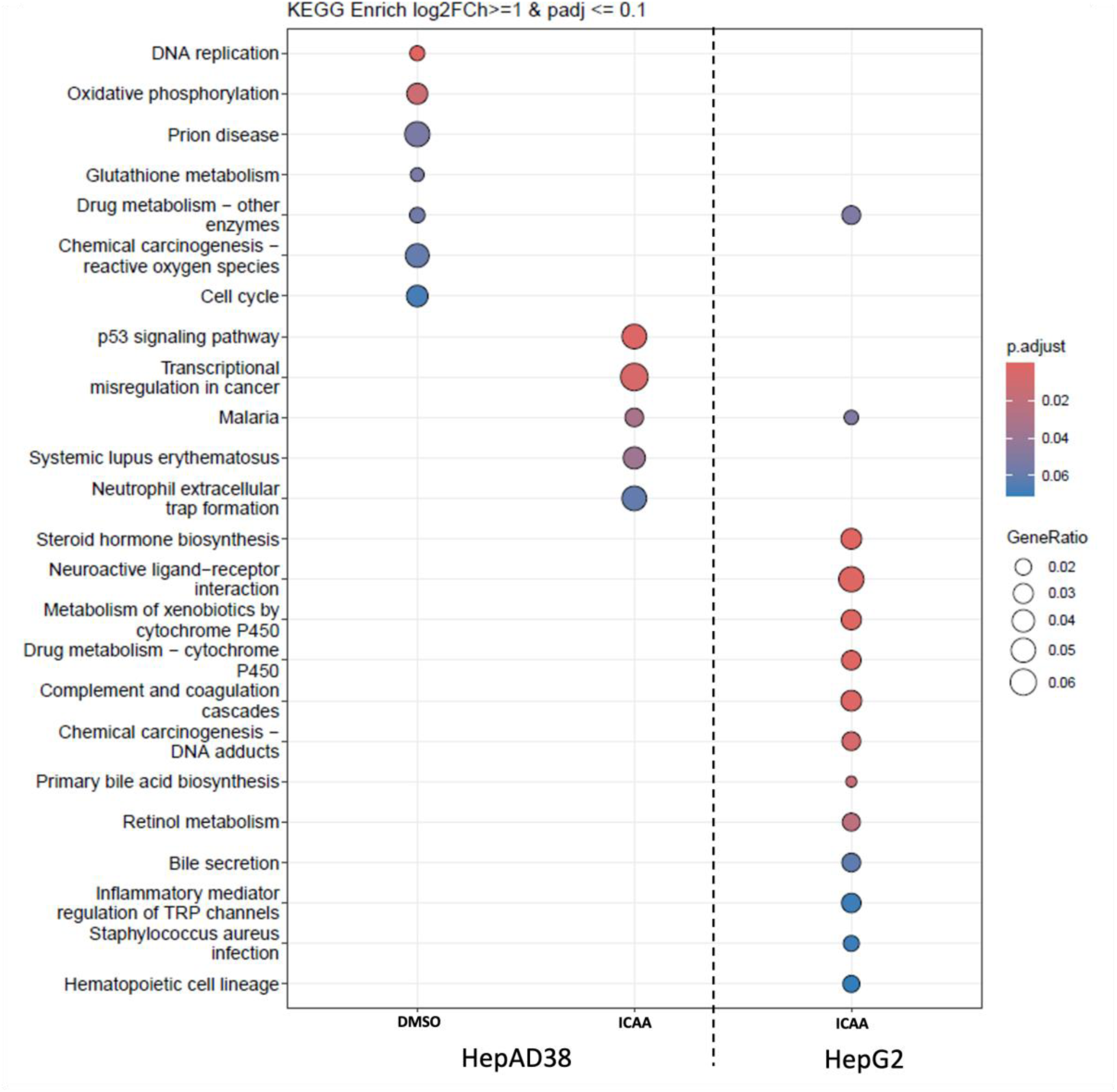
Top ranked KEGG enriched pathways of DEGs in HepG2 and HepAD38 3 days post-treatment. The first half of the graph illustrates significantly enriched KEGG pathways identified from DMSO- and ICAA-treated HepAD38, representing genes regulated by HBV-associated and ICAA-treated conditions in HepAD38. The second half corresponds to genes modulated following ICAA treatment in HepG2 cells. Each dot represents an enriched pathway, with color intensity corresponding to enrichment significance (p-value) and dot size proportional to the number of genes associated with each pathway.

**Supplemental figure 3.**
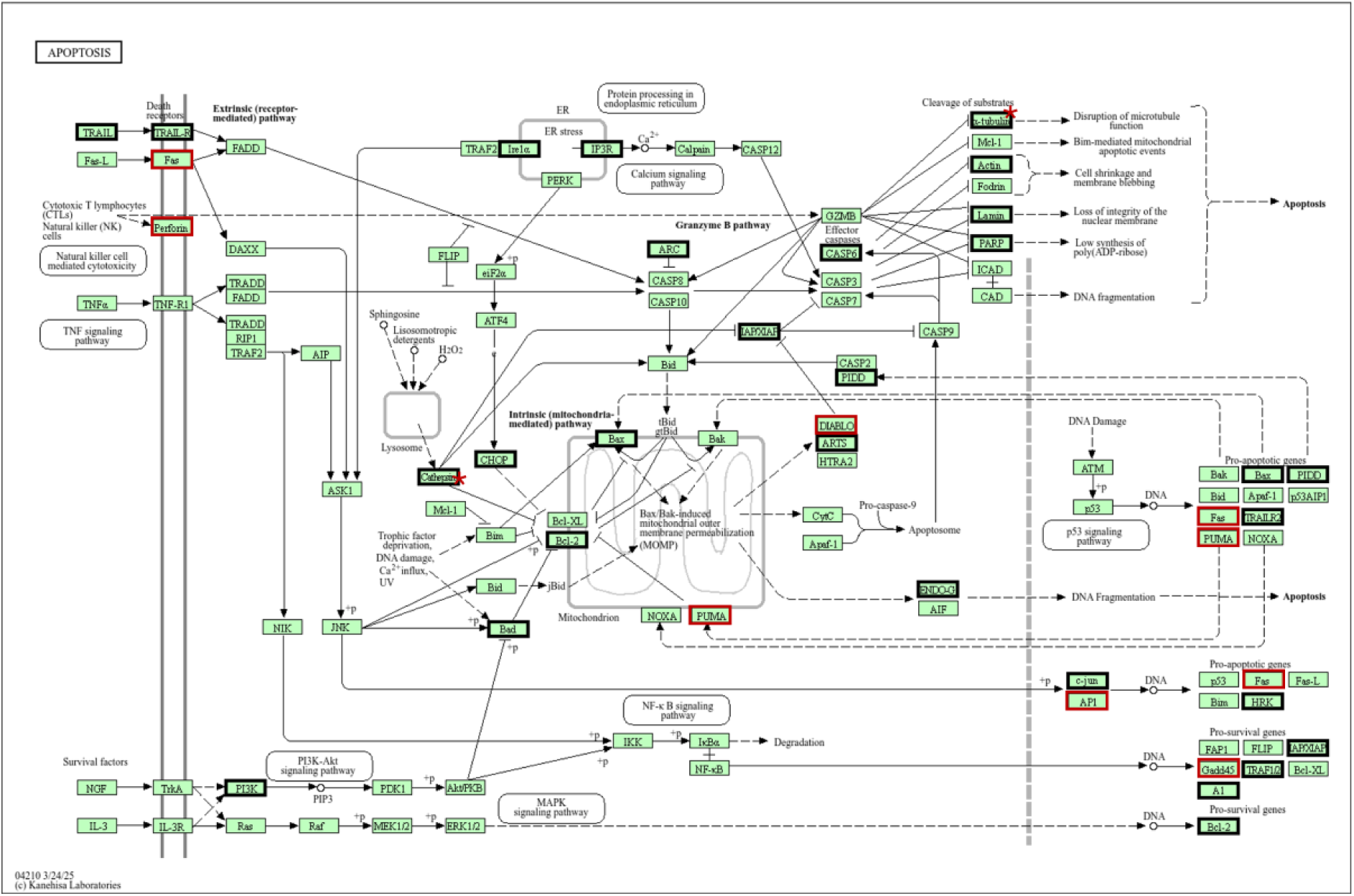
KEGG Pathview of the apoptosis signaling pathway. The Pathview illustrates the apoptosis activation process and its components. The dark blocks indicate the 28 components modulated by HBV, while the red blocks indicate the components modulated by ICAA. The red stars represent those identified to share common nodes between the two conditions.

## References

1. Tsukuda, Senko, and Koichi Watashi. 2020. Hepatitis B virus biology and life cycle. Antiviral Research 182. Elsevier: 104925. 10.1016/J.ANTIVIRAL.2020.104925.

2. Liang, T. Jake. 2009. Hepatitis B: The Virus and Disease. *Hepatology (Baltimore, Md.)* 49. NIH Public Access: S13. 10.1002/HEP.22881.

3. Nassal, Michael. 2015. HBV cccDNA: viral persistence reservoir and key obstacle for a cure of chronic hepatitis B. Gut 64. Gut: 1972–1984. 10.1136/GUTJNL-2015-309809.

4. Qi, Yonghe, Zhenchao Gao, Guangwei Xu, Bo Peng, Chenxuan Liu, Huan Yan, Qiyan Yao, et al. 2016. DNA Polymerase κ Is a Key Cellular Factor for the Formation of Covalently Closed Circular DNA of Hepatitis B Virus. PLOS Pathogens 12. Public Library of Science: e1005893. 10.1371/JOURNAL.PPAT.1005893.

5. KöNiger, Christian, Ida Wingert, Moritz Marsmann, Christine Rösler, Jörgen Beck, and Michael Nassal. 2014. Involvement of the host DNA-repair enzyme TDP2 in formation of the covalently closed circular DNA persistence reservoir of hepatitis B viruses. Proceedings of the National Academy of Sciences of the United States of America 111. Proc Natl Acad Sci U S A: E4244–E4253. 10.1073/PNAS.1409986111.

6. Easterbrook, Philippa J, Niklas Luhmann, Sahar Bajis, Myat Sandi Min, Morkor Newman, Olufunmilayo Lesi, and Meg C Doherty. 2024. WHO 2024 hepatitis B guidelines: an opportunity to transform care. 10.1016/S2468-1253(24)00089X.

7. Chien, Rong Nan, and Yun Fan Liaw. 2022. Current Trend in Antiviral Therapy for Chronic Hepatitis B. Viruses 14. Viruses. 10.3390/V14020434.

8. Tu, Thomas, Magdalena A. Budzinska, Nicholas A. Shackel, and Stephan Urban. 2017. HBV DNA Integration: Molecular Mechanisms and Clinical Implications. Viruses 9. Viruses. 10.3390/V9040075.

9. Yun, Chawon, Hae Ryun Um, Young Hee Jin, Jin Hee Wang, Mi Ock Lee, Sun Park, Jae Ho Lee, and Hyeseong Cho. 2002. NF-κB activation by hepatitis B virus X (HBx) protein shifts the cellular fate toward survival. Cancer Letters 184. Elsevier: 97–104. 10.1016/S0304-3835(02)00187-8.

10. Chao, Chuck C.K. 2016. Inhibition of apoptosis by oncogenic hepatitis B virus X protein: Implications for the treatment of hepatocellular carcinoma. World Journal of Hepatology 8. Baishideng Publishing Group Co: 1061. 10.4254/WJH.V8.I25.1061.

11. Quirino, Angela, Nadia Marascio, Francesco Branda, Alessandra Ciccozzi, Chiara Romano, Chiara Locci, Ilenia Azzena, et al. 2024. Viral Hepatitis: Host Immune Interaction, Pathogenesis and New Therapeutic Strategies. Pathogens 13. Multidisciplinary Digital Publishing Institute (MDPI): 766. 10.3390/PATHOGENS13090766.

12. Kuwata, Kazuo, Tomohiko Urushisaki, Tomoaki Takemura, Shigemi Tazawa, Mayuko Fukuoka, Junji Hosokawa-Muto, and Yoko Araki. 2011. Caffeoylquinic Acids Are Major Constituents with Potent Anti-Influenza Effects in Brazilian Green Propolis Water Extract. Evidence-based Complementary and Alternative Medicine: eCAM 2011: 254914. 10.1155/2011/254914.

13. Makola, Mpho M., Ian A. Dubery, Gerrit Koorsen, Paul A. Steenkamp, Mwadham M. Kabanda, Louis L. Du Preez, and Ntakadzeni E. Madala. 2016. The Effect of Geometrical Isomerism of 3,5-Dicaffeoylquinic Acid on Its Binding Affinity to HIV-Integrase Enzyme: A Molecular Docking Study. Evidence-Based Complementary and Alternative Medicine 2016. John Wiley & Sons, Ltd: 4138263. 10.1155/2016/4138263.

14. Koyaweda, Giscard Wilfried, Mirco Glitscher, Anja Schollmeier, Daniela Bender, and Eberhard Hildt. 2026. Isochlorogenic acid A impairs hepatitis B virus replication by interference with various steps of hepatitis B virus life cycle involving HO-1-mediated ROS modulation. Antiviral Research 245. Elsevier: 106323. 10.1016/J.ANTIVIRAL.2025.106323.

15. Guo, Jun Tao, Han Yu Li, Chao Cheng, Jia Xue Shi, Hai Nan Ruan, Jun Li, and Chan Min Liu. 2024. Lead-induced liver fibrosis and inflammation in mice by the AMPK/MAPKs/NF-κB and STAT3/TGF-β1/Smad2/3 pathways: the role of Isochlorogenic acid a. Toxicology Research 13. Oxford University Press. 10.1093/toxres/tfae072.

16. WANG, Jing, Hong WANG, Ying PENG, Guang Ji WANG, and Hai Ping HAO. 2016. Isochlorogenic acid A affects P450 and UGT enzymes in vitro and in vivo. Chinese journal of natural medicines 14. Chin J Nat Med: 865–870. 10.1016/S1875-5364(16)30103-0.

17. Kim, Joo Youn, Hong Kyu Lee, Bang Yeon Hwang, Seung Hwan Kim, Jae Kuk Yoo, and Yeon Hee Seong. 2012. Neuroprotection of ilex latifolia and caffeoylquinic acid derivatives against excitotoxic and hypoxic damage of cultured rat cortical neurons. Archives of Pharmacal Research 35. Springer: 1115–1122. 10.1007/S12272-012-0620-Y/METRICS.

18. Ladner, Stephanie K, Michael J Otto, Christopher S Barker, † Katie Zaifert, Guang-Hua Wang, Ju-Tao Guo, Christoph Seeger, and Robert W King. 1997. Inducible Expression of Human Hepatitis B Virus (HBV) in Stably Transfected Hepatoblastoma Cells: a Novel System for Screening Potential Inhibitors of HBV Replication. Vol. 41.

19. Levin, Joshua Z., Moran Yassour, Xian Adiconis, Chad Nusbaum, Dawn Anne Thompson, Nir Friedman, Andreas Gnirke, and Aviv Regev. 2010. Comprehensive comparative analysis of strand-specific RNA sequencing methods. Nature methods 7. Nature Publishing Group: 709. 10.1038/NMETH.1491.

20. Chen, Shifu, Yanqing Zhou, Yaru Chen, and Jia Gu. 2018. fastp: an ultra-fast all-in-one FASTQ preprocessor. Bioinformatics 34. Oxford University Press: i884. 10.1093/BIOINFORMATICS/BTY560.

21. Dobin, Alexander, Carrie A. Davis, Felix Schlesinger, Jorg Drenkow, Chris Zaleski, Sonali Jha, Philippe Batut, Mark Chaisson, and Thomas R. Gingeras. 2012. STAR: ultrafast universal RNA-seq aligner. Bioinformatics 29: 15. 10.1093/BIOINFORMATICS/BTS635.

22. Love, Michael I., Wolfgang Huber, and Simon Anders. 2014. Moderated estimation of fold change and dispersion for RNA-seq data with DESeq2. Genome Biology 15. BioMed Central Ltd.: 550. 10.1186/S13059-014-0550-8.

23. Kanehisa, Minoru, Miho Furumichi, Yoko Sato, Yuriko Matsuura, and Mari Ishiguro-Watanabe. 2024. KEGG: biological systems database as a model of the real world. Nucleic Acids Research 53. Oxford University Press: D672. 10.1093/NAR/GKAE909.

24. Yu, Guangchuang, Li Gen Wang, Yanyan Han, and Qing Yu He. 2012. clusterProfiler: an R Package for Comparing Biological Themes Among Gene Clusters. OMICS: a Journal of Integrative Biology 16: 284. 10.1089/OMI.2011.0118.

25. Kreß, Julia Katharina Charlotte, Christina Jessen, Anita Hufnagel, Werner Schmitz, Thamara Nishida Xavier da Silva, Ancély Ferreira dos Santos, Laura Mosteo, Colin R. Goding, José Pedro Friedmann Angeli, and Svenja Meierjohann. 2023. The integrated stress response effector ATF4 is an obligatory metabolic activator of NRF2. Cell Reports 42. Elsevier B.V.: 112724. 10.1016/j.celrep.2023.112724.

26. Chung, Young Min, See Hyoung Park, Wen Bin Tsai, Shih Ya Wang, Masa Aki Ikeda, Jonathan S. Berek, David J. Chen, and Mickey C.T. Hu. 2012. FOXO3 signalling links ATM to the p53 apoptotic pathway following DNA damage. Nature communications 3. Nature Publishing Group: 1000. 10.1038/NCOMMS2008.

27. Ryan, K. M., A. C. Phillips, and K. H. Vousden. 2001. Regulation and function of the p53 tumor suppressor protein. Current Opinion in Cell Biology 13. Elsevier Current Trends: 332–337. 10.1016/S0955-0674(00)00216-7.

28. Rawat, Siddhartha, and Michael J. Bouchard. 2014. The Hepatitis B Virus (HBV) HBx Protein Activates AKT To Simultaneously Regulate HBV Replication and Hepatocyte Survival. Journal of Virology 89. American Society for Microbiology: 999. 10.1128/JVI.02440-14.

29. Itskanov, Samuel, Katie M. Kuo, James C. Gumbart, and Eunyong Park. 2021. Stepwise gating of the Sec61 protein-conducting channel by Sec63 and Sec62. Nature Structural & Molecular Biology 2021 28:2 28. Nature Publishing Group: 162–172. 10.1038/s41594-020-00541-x.

30. Ye, Zhi Wei, Jie Zhang, Tiffany Ancrum, Yefim Manevich, Danyelle M. Townsend, and Kenneth D. Tew. 2017. Glutathione S-Transferase P-Mediated Protein S-Glutathionylation of Resident Endoplasmic Reticulum Proteins Influences Sensitivity to Drug-Induced Unfolded Protein Response. Antioxidants & redox signaling 26. Antioxid Redox Signal: 247–261. 10.1089/ARS.2015.6486.

31. Satoh, Tadashi, Takayasu Toshimori, Gengwei Yan, Takumi Yamaguchi, and Koichi Kato. 2016. Structural basis for two-step glucose trimming by glucosidase II involved in ER glycoprotein quality control. Scientific Reports 6. Nature Publishing Group: 20575. 10.1038/SREP20575.

32. Lei, Yingying, Hong Yu, Shaoxue Ding, Hui Liu, Chunyan Liu, and Rong Fu. 2024. Molecular mechanism of ATF6 in unfolded protein response and its role in disease. Heliyon 10. Elsevier: e25937. 10.1016/J.HELIYON.2024.E25937.

33. Russell, Christopher, and Scott M. Stagg. 2010. New insights into the structural mechanisms of the COPII coat. Traffic 11. John Wiley & Sons, Ltd: 303–310. 10.1111/J.1600-0854.2009.01026.X;PAGE:STRING:ARTICLE/CHAPTER.

34. Hu, Chen, Jing Yang, Ziping Qi, Hong Wu, Beilei Wang, Fengming Zou, Husheng Mei, Jing Liu, Wenchao Wang, and Qingsong Liu. 2022. Heat shock proteins: Biological functions, pathological roles, and therapeutic opportunities. MedComm 3. John Wiley and Sons Inc: e161. 10.1002/MCO2.161.

35. Levrero, Massimo, and Jessica Zucman-Rossi. 2016. Mechanisms of HBV-induced hepatocellular carcinoma. Journal of Hepatology 64. Elsevier: S84–S101. 10.1016/J.JHEP.2016.02.021.

36. Takaki, Akinobu, and Kazuhide Yamamoto. 2015. Control of oxidative stress in hepatocellular carcinoma: Helpful or harmful? World Journal of Hepatology 7. Baishideng Publishing Group Co: 968. 10.4254/WJH.V7.I7.968.

37. D’souza, Simmone, Keith C.K. Lau, Carla S. Coffin, and Trushar R. Patel. 2020. Molecular mechanisms of viral hepatitis induced hepatocellular carcinoma. World Journal of Gastroenterology 26. Baishideng Publishing Group Co: 5759. 10.3748/WJG.V26.I38.5759.

38. Hildt, Eberhard, Barbara Munz, Gesine Sa Her, Kurt Reifenberg, and Peter Hans Hofschneider. 2002. The PreS2 activator MHBst of hepatitis B virus activates c-raf-1/Erk2 signaling in transgenic mice. The EMBO Journal 2002 21:4 21. Springer: 525–535. 10.1093/EMBOJ/21.4.525.

39. Hong, Xupeng, Elena S. Kim, and Haitao Guo. 2017. Epigenetic Regulation of Hepatitis B Virus Covalently Closed Circular DNA: Implications for Epigenetic Therapy against Chronic Hepatitis B. *Hepatology (Baltimore*, Md.) 66. John Wiley and Sons Inc.: 2066. 10.1002/HEP.29479.

40. Naully, Patricia Gita, Marselina Irasonia Tan, Agustiningsih Agustiningsih, Caecilia Sukowati, and Ernawati Arifin Giri-Rachman. 2025. cccDNA epigenetic regulator as target for therapeutical vaccine development against hepatitis B. Annals of Hepatology 30. Elsevier Espana S.L.U. 10.1016/j.aohep.2024.101533.

41. Rivière, Lise, Laetitia Gerossier, Aurélie Ducroux, Sarah Dion, Qiang Deng, Marie Louise Michel, Marie Annick Buendia, Olivier Hantz, and Christine Neuveut. 2015. HBx relieves chromatin-mediated transcriptional repression of hepatitis B viral cccDNA involving SETDB1 histone methyltransferase. Journal of Hepatology 63. Elsevier: 1093–1102. 10.1016/J.JHEP.2015.06.023.

42. Nguyen-Lefebvre, Anh Thu, Nazia Selzner, Jeffrey L. Wrana, and Mamatha Bhat. 2021. The hippo pathway: A master regulator of liver metabolism, regeneration, and disease. FASEB Journal 35. John Wiley and Sons Inc: e21570. 10.1096/FJ.202002284RR;REQUESTEDJOURNAL:JOURNAL:15306 860;WGROUP:STRING:PUBLICATION.

43. Liu, Yuchen, Xiaohui Wang, and Yingzi Yang. 2020. Hepatic Hippo signaling inhibits development of hepatocellular carcinoma. Clinical and Molecular Hepatology 26. The Korean Association for the Study of the Liver: 742–750. 10.3350/CMH.2020.0178.

44. Lin, Shaoli, and Yan Jin Zhang. 2017. Interference of Apoptosis by Hepatitis B Virus. Viruses 9. MDPI AG: 230. 10.3390/V9080230.

45. Liu, Wei, Teng Fei Guo, Zhen Tang Jing, and Qiao Yun Tong. 2019. Repression of Death Receptor–Mediated Apoptosis of Hepatocytes by Hepatitis B Virus e Antigen. The American Journal of Pathology 189. Elsevier: 2181–2195. 10.1016/J.AJPATH.2019.07.014.

46. Jabbar Yasir, Saif. 2026. HBV Infection and Apoptosis: Molecular Mechanisms and Clinical Implications. International Journal of Environmental and Biological Sciences 1. 10.63939/2CCAG975.

47. Schollmeier, Anja, Michael Basic, Mirco Glitscher, and Eberhard Hildt. 2024. The impact of HBx protein on mitochondrial dynamics and associated signaling pathways strongly depends on the hepatitis B virus genotype. Journal of Virology 98. American Society for Microbiology. 10.1128/JVI.00424-24/ASSET/8FDD3C9F-C8EB-4709-982A-2AD8A12F665A/ASSETS/IMAGES/LARGE/JVI.00424-24.F009.JPG.

48. Kim, Seong Jun, Mohsin Khan, Jun Quan, Andreas Till, Suresh Subramani, and Aleem Siddiqui. 2013. Hepatitis B Virus Disrupts Mitochondrial Dynamics: Induces Fission and Mitophagy to Attenuate Apoptosis. PLOS Pathogens 9. Public Library of Science: e1003722. 10.1371/JOURNAL.PPAT.1003722.

49. Xu, Jian Guo, Qing Ping Hu, and Yu Liu. 2012. Antioxidant and DNA-Protective Activities of Chlorogenic Acid Isomers. Journal of Agricultural and Food Chemistry 60. American Chemical Society: 11625–11630. 10.1021/JF303771S.

50. Kannan, Krishnaswamy, and Sushil K. Jain. 2000. Oxidative stress and apoptosis. Pathophysiology 7. Pathophysiology: 153–163. 10.1016/S0928-4680(00)00053-5.

51. Hockenbery, David M., Zoltan N. Oltvai, Xiao Ming Yin, Curt L. Milliman, and Stanley J. Korsmeyer. 1993. Bcl-2 functions in an antioxidant pathway to prevent apoptosis. Cell 75. Cell Press: 241–251. 10.1016/0092-8674(93)80066-N.

52. Chen, Yanchi, Yiling Li, Linyang Huang, Yu Du, Feihong Gan, Yanxi Li, and Yang Yao. 2021. Antioxidative Stress: Inhibiting Reactive Oxygen Species Production as a Cause of Radioresistance and Chemoresistance. Oxidative Medicine and Cellular Longevity 2021. Hindawi Limited: 6620306. 10.1155/2021/6620306.

